# Elementary Growth Modes provide a molecular description of cellular self-fabrication

**DOI:** 10.1101/608083

**Authors:** Daan H. de Groot, Josephus Hulshof, Bas Teusink, Frank J. Bruggeman, Robert Planqué

**Affiliations:** Systems Bioinformatics, Amsterdam Institute for Molecules, Medicines & Systems, VU University, De Boelelaan 1087, 1081 HV, Amsterdam, The Netherlands; Department of Mathematics, VU University, De Boelelaan 1081a, 1081 HV, Amsterdam, The Netherlands

## Abstract

A major aim of biology is to predict phenotype from genotype. Here we ask if we can describe all possible molecular states (phenotypes) for a cell that fabricates itself at a constant rate, given its enzyme kinetics and the stoichiometry of all reactions (the genotype). For this, we must understand the autocatalytic process of cellular growth which is inherently nonlinear: steady-state self-fabrication requires a cell to synthesize all of its components, including metabolites, enzymes and ribosomes, in the proportions that exactly match its own composition – the growth demand thus depends on the cellular composition. Simultaneously, the concentrations of these components should be tuned to accomplish this synthesis task – the cellular composition thus depends on the growth demand. We here derive a theory that describes all phenotypes that solve this circular problem; the basic equations show how the concentrations of all cellular components and reaction rates must be balanced to get a constant self-fabrication rate. All phenotypes can be described as a combination of one or more minimal building blocks, which we call Elementary Growth Modes (EGMs). EGMs can be used as the theoretical basis for all models that explicitly model self-fabrication, such as the currently popular Metabolism and Expression models. We then used our theory to make concrete biological predictions: we find that natural selection for maximal growth rate drives microorganisms to states of minimal phenotypic complexity: only one EGM will be active when cellular growth rate is maximised. The phenotype of a cell is only extended with one more EGM whenever growth becomes limited by an additional biophysical constraint, such as a limited solvent capacity of a cellular compartment. Our theory starts from basic biochemical and evolutionary considerations, and describes unicellular life, both in growth-promoting and in stress-inducing environments, in terms of EGMs, the universal building blocks of self-fabrication and a cell’s phenotype.

## 1 Introduction

One of the defining aspects of living cells is that they fabricate themselves from simple chemical compounds, a process often referred to as cell growth or replication. Microorganisms can do this in a multitude of different environments, using various food sources and while facing various stresses. Understanding the molecular players and mechanisms of self-replication is the realm of (molecular) microbiology; understanding the design principles underneath the regulatory adaptation processes, and how phenotype emerges from the molecular level, is the key question in systems biology. In evolutionary biology, the impact of phenotypic strategies on fitness is evaluated. Connecting these fields, and providing what is sometimes called the genotype-to-phenotype map, is a grand challenge in biology.

The aim of this paper is to make a start with just that: we provide a molecular mechanistic description of how a cell maintains itself and how it grows, in terms of all its biosynthetic processes and stress-managing systems. This may seem an impossible task, but when we restrict ourselves to steady state exponential, or balanced, growth (Fishov et al., 1995), this becomes a feasible endeavour, as we will show. In such a balanced growth state, all intrinsic properties of a population of cells, such as distributions of molecule concentrations and reaction rates, are time-invariant. Under these circumstances, a molecular mechanistic description of a cellular phenotype amounts to listing all reaction rates and all the concentrations of cellular components.

Arguably the most important phenotypic trait of growing microbial cells is their growth rate, since it directly determines the number of offspring cells synthesised per unit time. In growth-promoting conditions, the vast majority of a cell’s metabolic energy and resources is invested into growth and the production of new cells. There is also a direct evolutionary premium on fast growth: under constant conditions, specific growth rate is the direct determinant of fitness. We are therefore interested in understanding how a certain cellular phenotype gives rise to its steady state exponential growth rate.

To control the growth rate, proteins and ribosomes need to be expressed at different concentrations; to attain higher growth rates, the rates of enzyme synthesis need to be higher, both to achieve higher enzyme concentrations and thus higher reaction rates, but also to counter higher dilution rates by cell growth. Higher enzyme synthesis rates require higher numbers of ribosomes, but since the ribosomes also produce these additional ribosomes, we need even more ribosomes. The relative abundance of ribosomes and enzymes will therefore inevitably change with increasing growth rate, turning cellular growth into a nonlinear problem. This problem is solved by exactly balancing the synthesis rates of all enzymes and the ribosome. Therefore, the control space of the cell can ultimately be viewed as a ribosomal allocation space, with different fractions of the ribosome allocated to synthesise the different enzymes. These fractions might be determined by passive competition for mRNA molecules produced by gene expression. The aim of this paper is to classify all possible balanced growth states of a cell in terms of those ribosomal fraction variables.

There is a long tradition in trying to model growth of (microbial) cells, but in particular genomescale metabolic network models have come close to providing a comprehensive molecular description of cellular phenotypes (Price et al., 2004). These models link genes to metabolic enzymes to reaction stoichiometries, and the reactions form a metabolic network through which matter flows in the form of metabolic flux from sources (nutrients) to sinks (biomass and byproducts). Such models neglect enzyme kinetics and the synthesis of enzymes and ribosomes, which greatly facilitates the analysis.

In order to predict growth phenotypes in such models, growth rate is assumed to be proportional to the rate of a so-called biomass reaction: a virtual reaction with fixed stoichiometry that is used to model the average demand for biomass components. This biomass reaction rate is being maximised at steady state under governing constraints on fluxes—usually input fluxes. This constitutes a linear optimisation problem with the individual fluxes as the optimisation (or control) variables, and is well known as Flux Balance Analysis (FBA; Orth et al. (2010)). We understand the solution space of an FBA problem very well from a mathematical point of view (Schuster et al., 2000; Gagneur and Klamt, 2004; Bordel and Nielsen, 2010). This understanding was facilitated by the identification of a set of invariant building blocks of the solution space: the Elementary Flux Modes (EFMs) (Schuster and Hilgetag, 1994; Schuster et al., 2002).

In recent years, cellular resource management has become an important concept with which to improve our understanding of cellular physiology (Molenaar et al., 2009; Scott et al., 2010; Weiße et al., 2015; Basan et al., 2015), even hinting at true ‘bacterial growth laws’ (Scott and Hwa, 2011). The additional insight offered by cellular resource management has been driving the field from classical FBA to “resource”-FBA type approaches. In these approaches a virtual biomass reaction is still optimized, but by introducing enzyme kinetics (either by introducing the full nonlinear enzyme saturation function, or by introducing only a maximal rate (k_cat_) per enzyme) the constraints now act on the enzyme concentrations, instead of on the fluxes. This creates an optimisation problem with the enzyme concentrations as the optimisation variables, a problem that is nonlinear when nonlinear enzyme kinetics were introduced. The solution space, and the optimal solutions, can still be understood in terms of EFMs (Wortel et al., 2014; Müller et al., 2014; de Groot et al., 2019).

The concepts of cellular resource management have been further exploited in a number of modelling approaches in which metabolic flux is explicitly coupled to the synthesis of enzyme, which in turn is coupled to the presence of ribosomes and its required substrates (amino acids and a source of free energy). In these approaches the demand for biomass components is a variable that is calculated in the model, rather than a fixed biomass reaction. So far, such models, be it core models that include kinetics (Molenaar et al., 2009; Weiße et al., 2015), or augmented genomescale metabolic and expression (ME) models (Goelzer et al., 2009; Lerman et al., 2012; O’Brien et al., 2016), have been used for simulation studies and produce biologically relevant regulatory behaviours, such as overflow metabolism (Molenaar et al., 2009) or catabolite repression (You et al., 2013). However, they differ in the nature of the imposed constraints, and still ignore (different) parts of the self-fabrication process. More importantly perhaps, there is a lack of understanding of the mathematical structure of the solutions to all these optimal-resource allocation problems—understanding we do have for FBA-type problems.

In this paper, we develop theory for this next generation of growth models. Our theory identifies a set of invariant building blocks for the steady-state solution space of a self-fabricating biochemical system, including all metabolites, enzymes, ribosomes, and their synthesis rates. We have called them Elementary Growth Modes, as they play a similar—but not exactly the same—role as EFMs in the metabolic network optimisation problem.

The structure of this paper is as follows. After introducing the relevant notation, we focus on a population of growing cells in balanced growth, and start by giving a definition of the growth rate in terms of metabolic rates inside these cells. Then we introduce the class of whole-cell models studied in this paper, and derive a set of relations that have to hold in balanced growth and which feature the control variables of the cell most transparently. The main remaining aim of the paper is then to study the solution space of these relations in terms of these control variables.

We first introduce Elementary Growth States (EGSs) and show that all balanced growth states, at a given growth rate and a fixed set of metabolite concentrations, can be formed by taking suitable convex combinations of such states. EGSs with the same participating reactions, but at different growth rates and metabolite concentrations, may be identified with each other, and such a class of EGSs is termed an Elementary Growth Mode (EGM). We then show that, if no additional biophysical constraints are introduced, balanced growth rate is maximised in exactly one such EGM. If multiple biophysical constraints are active, a mixture of EGSs may arise as the growth-rate maximiser.

Finally, it is a natural question to ask how the new EGSs and EGMs relate to the older EFMs. We show that under one additional biological assumption, each EGM can be mapped to a unique growth-sustaining EFM. We indicate using experimental data that this biological assumption is indeed borne out.

At the end of each section, we give a brief description of the biological interpretation and consequences of the results derived. We have tried to keep these separate from the mathematical theory, to aid both the more mathematically and the more biologically inclined reader.

## 2 Methods and results

**Notation** Vectors are denoted by boldface, e.g., ***v***. The vector ***u***_*j*_ denotes the *j*-th elementary vector, filled with zeros except for the *j*-th element, which is 1. The dot product between vectors ***v*** and ***w*** is denoted by ***v · w***, and |***v***| denotes Σ_*i*_|*v_i_*|, which in this paper always simplifies to Σ_*i*_*v_i_* because the vectors in question have positive entries. Time derivatives are denoted by overdots, e.g., 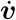. The inequality ***v*** ≥ 0 should be interpreted as *v_j_* ≥ 0 for all *j*. Modelling assumptions are labelled as A1, A2, etc. An overview of all variables and parameters used in this paper can be found in the Supplementary Information Table **??**.

### 2.1 Linking metabolic activity to the growth rate

Let ***n***(*t*) = (*n*_1_(*t*),…,*n_K_*(*t*)) be the vector of copy numbers of *K* compounds in a population of cells with total volume *V*(*t*) at time *t*. The concentration of compound *k* is defined as *c_k_*(*t*):= *n_k_*(*t*)/*V*(*t*). The modelling assumption is that changes in copy numbers of the compounds are due to chemical reactions which take place in a well-stirred cell(Heinrich and Schuster, 1996), so that

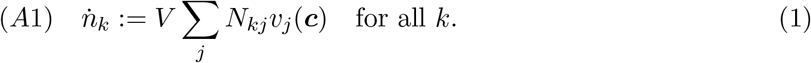

Here *N* denotes the stoichiometry matrix, and *v_j_*(***c***) is the *j*-th reaction rate, as a function of concentrations ***c***. Since *n_k_* = *Vc_k_*, we have

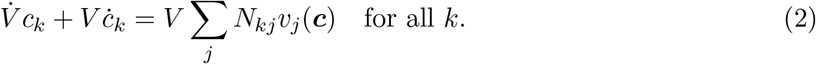

In balanced growth, all concentrations are constant over time (Schaechter, 2015), *ċ* = 0, which means that

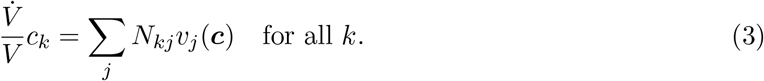

To relate the growth rate of the volume of a cell to all the metabolic reactions making new cell material, we have to relate cell volume to the molecule copy number. We consider experimental conditions such that the volume is a function of all the copy numbers only, *V*(*t*) = *V*(***n***(*t*)). We will discuss the implicit assumptions here at the end of this section. The time derivative of *V* = *V*(***n***) is given by

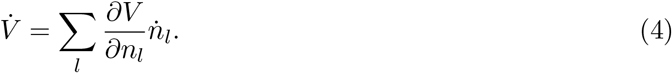

Hence, using (4) and (1) in (3), we have in balanced growth that

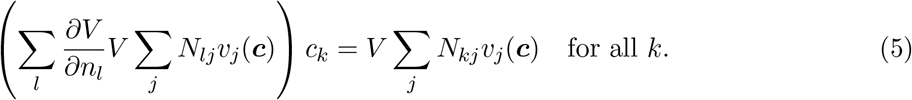

The steady-state assumption ensures that the concentrations ***c***, and therefore also the reaction rates *v_j_*(***c***) are constant in time. Therefore, the complete right-hand side of (5) is constant in time, and thus the left-hand side must be as well. This strongly suggests (but does not directly imply) that 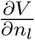 is independent of time as well.^1^ Since ***n*** does change in time, this implies that 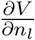 does not depend explicitly on ***n***, i.e.:

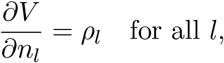

for constants *ρ_l_*.^2^ The biophysically reasonable assumption that total volume is the sum of all the volumes taken up by individual molecules,

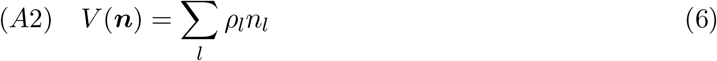

is therefore suitable to study balanced growth states. With this definition, we also have

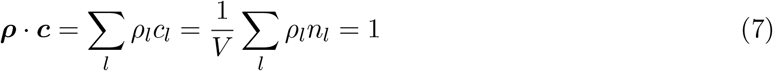

at balanced growth.

The (instantaneous) growth rate is now defined as

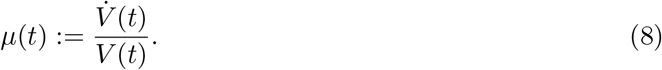

Since ***ρ · c*** = 1 at balanced growth,

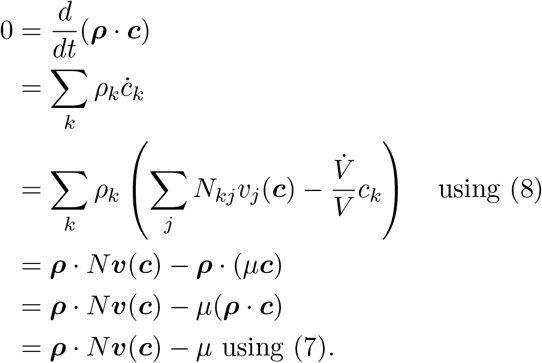

We conclude that the growth rate of a cell at balanced growth can be expressed in terms of the catalytic activities of its constituents reactions as,

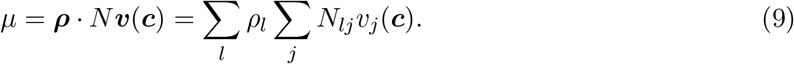

Since *ρ_l_* equals the molar volume of molecule *l*, this last relation simply states that the growth rate of a cell equals the total volume synthesis rate of all its reactions per unit volume. This total volume synthesis rate is calculated by summing the volumes of all synthesized metabolites in all reactions per unit time.

Equation (2) is equivalent to

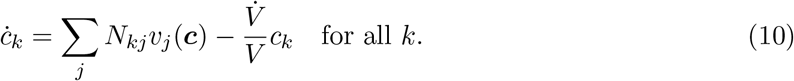

If we substitute the balanced growth rate (9) into (10) at steady state conditions, we obtain

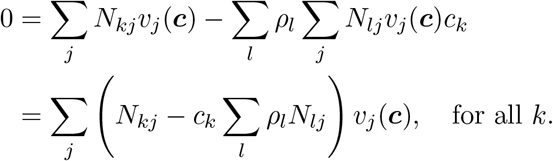

This equation shows that at steady-state, the net production of molecule *k* (first term) should balance the dilution of molecule *k* by cellular growth, growth that is due to the volume contribution of all produced molecules in the cell (second term). We could not analyse these equations further to understand the structure of steady-state solutions, specifying the metabolic activities of a cell that grows balanced, without additional mathematical assumptions.

#### Biological interpretation and consequences

In the previous section we related the growth rate of a population of cells to the volume changes due to the activities of metabolic reactions. As such, we showed that growth is due to the production of cell components, such as DNA, RNA, lipids and proteins, from extracellular nutrients. Our definition of growth rate is closely related to how growth rate is measured, using for instance optical density measurements.

In our derivation, we assumed that *V*(*t*) = *V*(***n***(*t*)), which means that the contents of a cell is approximated by an ideal solution (Dill and Bromberg, 2012). We realise that this is likely an oversimplifying assumption (McGuffee and Elcock, 2010; Parry et al., 2014)), but one that underlies many models of cell growth (de Jong et al., 2017; Schaechter, 2015). We note that balanced growth becomes much harder to rationalise when the assumption *V*(*t*) = Σ_*k*_ *ρ_k_n_k_*(*t*) is invalid. Therefore we have effectively assumed that cells grow at constant osmotic pressure. We remark that it is indeed likely that cells maintain a constant osmotic pressure during growth and that changes in extracellular osmotic pressure are compensated for by volume changes of the cell.

The theory so far concerns the descriptions of the copy number and volume changes in a population of cells that grows balanced, i.e., at a fixed growth rate, and such that all extrinsic properties increase exponentially and all intensive properties are constant (Schaechter, 2015). The relation between the properties of a population of cells to those of its individual cell members can be found in (Painter and Marr, 1968).

### 2.2 Introducing a whole-cell model of self-fabricating cells

To obtain a better understanding of how the molecular state of a cell, its ‘phenotype’, relates to its growth rate, we have to add more biological information to the general model from Section 2.1.

We distinguish concentrations of metabolites ***x***, enzymes ***e*** and ribosomes *r*. We ignore several biological entities, especially mRNA and genes. Genes are implicitly defined by the reaction stoichiometries occurring in the stoichiometric matrix; they are not synthesized nor degraded. DNA itself can be treated as a macromolecule in our description. Messenger RNA is not considered in our description as an entity but does play a role, as will become clear later. We note that the model can be extended with mRNA, without qualitative changes. We left it out for simplicity and because, in terms of volume, it is a minor cellular component, in contrast to rRNA, which can be considered as a regular macromolecule.

We note that in all population perspectives on cell growth, including ours, the division and birth process is generally not incorporated as a molecular process. All cells are growing and dividing asynchronously, and are considered in the same state of producing cell components. At each moment in time, a fixed fraction of cells is dividing (Painter and Marr, 1968) in a population of cells at balanced growth; considering their activities would require a different modelling formalism than the one pursued in this paper. We have chosen this formalism because it is the overarching description of all genome-scale modelling formalisms (Price et al., 2004; O’Brien et al., 2016) that are currently in use in systems biology.

The dynamical system for all concentrations ***c*** = (***x, e***, *r*) is still given by (10). We subdivide the stoichiometric matrix *N* into parts, corresponding to the two levels of “metabolism” and “enzymes and ribosome synthesis”, as follows,

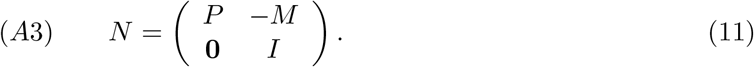

Here *P* is an *m* × *n*-matrix (the usual stoichiometric matrix in a metabolic pathway), with entries generally of different signs. The number of metabolites needed for the enzyme synthesis reactions are given by *M*, an *m* × (*n* + 1)-matrix with mostly positive (or zero) entries. The only negative entries in *M* correspond to metabolites that are produced when enzymes are synthesized, such as ADP. The (*n* + 1) × (*n* + 1) identity matrix, *I*, denotes that each enzyme is made by one enzyme synthesis reaction. The vector of reaction rates is also split, into metabolic rates ***v*** = (*v*_1_,…,*v_n_*), and synthesis reactions ***w***, consisting of *n* enzyme synthesis rates (*w*_1_,…*w_n_*) and a ribosome synthesis rate *w*_*n*+1_.

The assumptions underlying (11) are that each individual metabolic reaction *v_j_*(***c***) has a unique enzyme associated to it. Since enzyme *j* catalyses metabolic reaction *j*, we assume, in agreement with basic enzyme kinetics (Cornish-Bowden, 2004), that

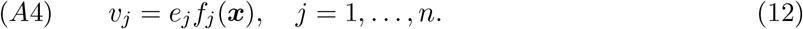

The ribosome catalyses synthesis of all enzymes and itself, and amino acids (which form part of the metabolites ***x***) are consumed in the process. We therefore choose

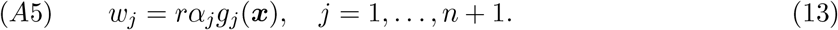

The rate laws *f_j_*(***x***) are assumed to be known nonlinear functions, and contain the catalytic rate parameter *k*_cat,*j*_ for the enzyme *e_j_* in this notation; moreover, they incorporate the thermodynamic constraints imposed by the substrates and products of each enzyme. Also the *g_j_*(***x***) are assumed to be known and contain *k_cat,r_*. The linear dependence of the enzymatic rate laws *f_j_* follows generally from quasi-steady-state type derivations of rate laws (Cornish-Bowden, 2004), and are a good approximation when enzymes are at concentrations that are much lower than the substrate or product concentrations. The linear dependence of enzyme and ribosome synthesis rates on ribosome level in (13) is assumed for the same reason. Both linear dependencies are crucial for the development of the theory.

The coefficient *α_j_* is the fraction of the total ribosome pool that is allocated to produce enzyme *j*, and are the result of gene expression. A possible mechanism is that the different mRNA molecules passively compete to be translated by the ribosome. The fraction of the ribosome allocated to produce enzyme *j* is then assumed to be proportional to the fraction of mRNA_*j*_ over total mRNA (but not necessarily equal to it, since a certain fraction of the ribosome may not be allocated to anything, in principle). Our theory does not depend on the specific mechanism that is used in cells, but is based solely on the assumption that the *α_j_*-factors can be influenced by gene expression. We have

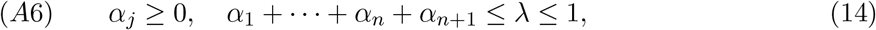

where λ signifies the fraction of all the ribosomes that are actively involved in synthesis of new protein and ribosomes. We emphasise that the *α_j_*’s are the main control parameters of the cell in the perspective taken in this paper. By changing them, the cell changes phenotype.

Let us lastly introduce the molar volume parameters for the compounds. We denote these by *ρ*_1_,…,*ρ_m_* for the *m* metabolites, *σ*_1_,…,*σ_n_* for the *n* enzymes, and *σ*_*n*+1_ for the ribosome. Assumption (A2) now reads, analogously to (7),

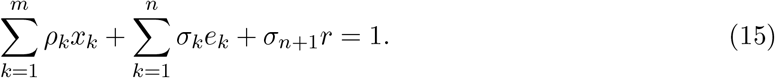

In summary, assumptions (A1)–(A6) give the following dynamical system for a whole-cell model,

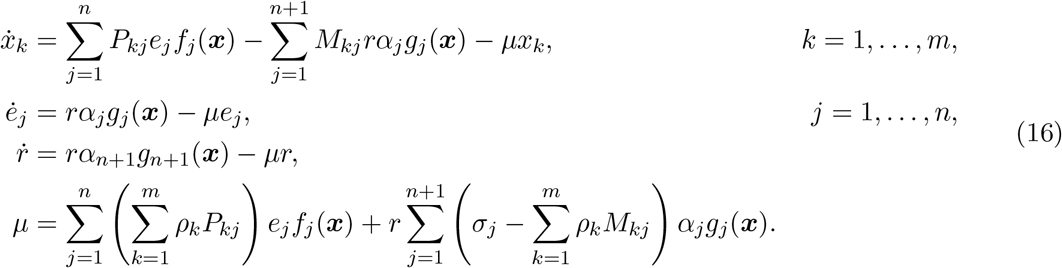

External environmental concentrations, such as nutrients, are incorporated as parameters in the relevant *f_j_*(***x***).

#### Biological interpretation and consequences

The growth rate given in (16) has a natural biological interpretation. The overall growth rate of cells is a sum of the net relative volume increases due to metabolic reactions (including transport reactions, and internal reactions in which volumes of substrate molecules are replaced by volumes of product molecules), and the net volume change as the result of the conversion of metabolites into proteins and ribosomes. The quantity 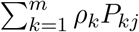 is the net change in volume due to the conversion of substrates into products in metabolic reaction *j*, and 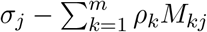 is the net volume change due to the production of one mole of enzyme j from the relevant amino acids. These quantities will be called *a_j_* and *b_j_* below. For different choices of ***α***, different sets of enzymes are synthesised resulting in different growth rates.

Note that the dependence of *μ* on the concentrations (*<u>x</u>*, *<u>e</u>*, *r*) forms an important aspect of our theory. If only the first three equations of (16) were to be solved, then we would model the behaviour of a set of concentrations in an exponentially growing volume. The expression for *μ* however indicates that the growth rate of this volume is dependent on this same set of concentrations. Cellular growth is thus really coupled to the chemical reactions in the cell.

### 2.3 The balanced growth equations

To make notation more concise, set

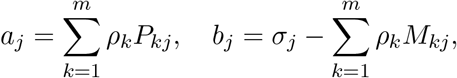

so that *μ* in (16) simplifies to

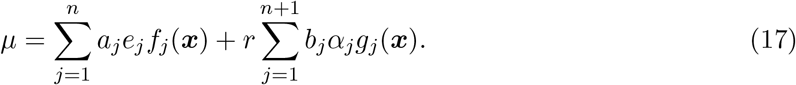

We now derive a set of identities that need to hold at balanced growth, i.e., when ***ċ*** = 0. The aim is that they are written in such as a way that the variables under control of the cell, the ribosomal allocation parameters *α_j_*, stand out.

Since *ė_j_* = 0, *μe_j_* = *rα_j_g_j_*(***x***), and the definition of *μ* in (17) may be rewritten as the following equation for *μ*,

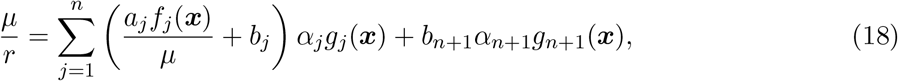

which allows us to compute a steady state value for *r* in terms of ***x***, *α*_1_,…,*α*_*n*+1_ and *μ*,

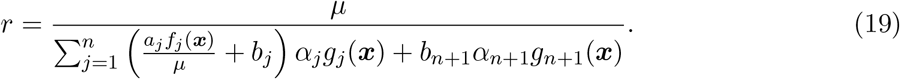

Demanding *x̄_k_* = 0 for *k* = 1,…,*m* in Equation (16) gives another *m* identities for *μ*,

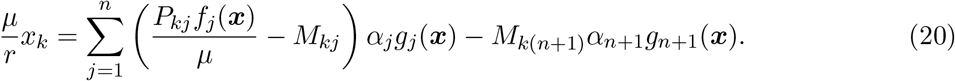

With Equations (18) and (20), we now have two different expressions of 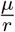 that should be equal whenever *x_k_* ≠ 0. This yields

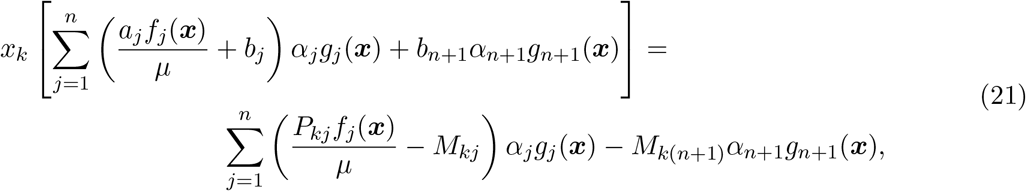

for *k* = 1,…,*m*, which are *m* equations for ***x*** and *μ* only.

Lastly, we still need to require *ṙ* = 0, which yields

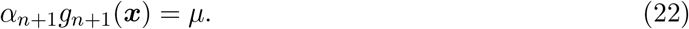

It is tempting to use this relation straight away in (21), but it is much better not to submit to this temptation as it would destroy the geometric structure of the solution space in the remaining fractions *α*_1_,…,*α_n_*. This is a vital ingredient later in the construction of Elementary Growth States and Modes.

For future reference, we collect the pertinent equations (21), (22), together with the assumptions on ***α*** in (14), and collectively call these the Balanced Growth equations,

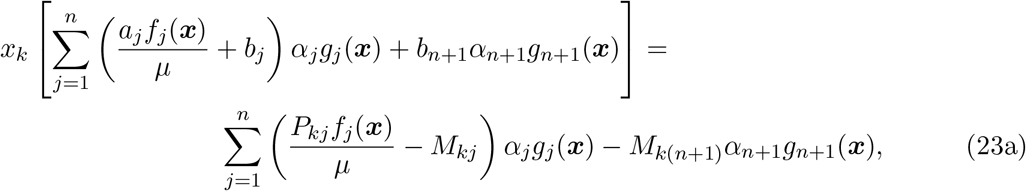

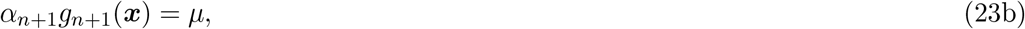

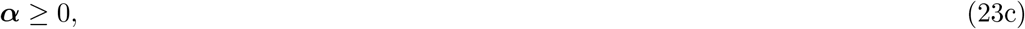

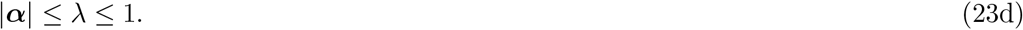

Note that when a specific ribosome allocation is chosen (by fixing ***α***), we get *m* + 1 equations that should be met. In principle, this is enough to fix the *m* metabolite concentrations and the growth rate, *μ*. Subsequently, the enzyme and ribosome concentrations, ***e***, *r*, can also be calculated. The *α_j_*-factors can thus be seen as controlling all other variable quantities in the model. However, not all choices for ***α*** will yield a solvable system of equations, and not all ribosome allocations will thus lead to balanced growth.

#### Biological interpretation and consequences

The phenotypic space of cells growing at a steady state rate consists of all possible sets of molecule concentrations that admit balanced growth. Each environmental condition, together with a particular ribosome allocation that allows growth, yields one phenotypic state. The complete phenotypic space is typically very large: many cells are able to attain balanced growth in many different conditions, often using different metabolic subnetworks (Kelk et al., 2012). Even on the same food substrates, metabolism may change when the concentrations of these substrates change, e.g., from pure respiration to respirofermentation. The cells can adapt to conditions by changing enzyme concentrations through gene expression control.

The control space of the cell is thus in terms of the proteins that are being expressed; these are the variables that allow phenotypic changes, and the ones that need to be tuned to maximise fitness. We have rewritten the steady-state equations of the whole-cell model to allow an investigation of the solution space in terms of these control variables, the *α*’s. In other words, we can now start to explore the possible phenotypes a cell may express in terms of the biochemical properties (encoded in rate laws *f_j_* and *g_j_*) of the catalysing molecules, the enzymes and ribosomes, and the extracellular nutrient concentrations.

### 2.4 Some basic Linear Programming

We give a short review of some basic Linear Programming (see e.g. (Matoušek and Gärtner, 2007; Bertsimas and Tsitsiklis, 1997)) necessary for the definition of Elementary Growth States and Elementary Growth Modes.

Let *A* be an *m × n* matrix, *m* < *n*. The linear programs in this paper are all in *equational* or *standard form*

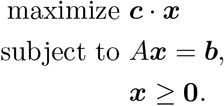

We may assume that *A* has full rank *m* (if *A* does not have full rank the matrix may be reduced to a matrix with fewer rows which does have full rank without changing the solution space of vectors satisfying *A****x*** = **b**). A vector ***x*** is called a *feasible solution* if it satisfies all the constraints, so *A****x*** = ***b*** and ***x*** ≥ **0**.

Let *D* ⊆ {1,…,*n*} be an index set, and let *A_D_* denote the matrix consisting of columns from *A* whose indices are in *D*. We will use similar notation for the restriction of ***x*** to the index set *D*, ***x***_*D*_. A vector ***x*** is called a *basic feasible solution* if ***x*** is a feasible solution for which there exists a set *D* ⊆ {1,…,*n*} with m distinct elements such that *A_D_* is square and non-singular, ***x*** satisfies *A_D_****x***_*D*_ = ***b***, and *x_j_* = 0 for all *j* ∉ *D*. The vector ***x***_*D*_ is then uniquely defined, since 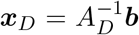. For any index set *D* with *A_D_* invertible, we can compute ***x***_*D*_. If this solution satisfies ***x***_*D*_ ≥ 0, then it induces a basic feasible solution ***x*** that satisfies *x_j_* = 0 for all indices of *j* ∉ *D. D* is then called a feasible basis for ***x***. (This parlance is different from the usual definition of a basis in Linear Algebra, in which a basis is formed by a set of vectors.)

We also recall a few standard concepts from convex geometry. A set *X* ⊂ ℝ^*n*^ is called *convex* if for each ***x, y*** ∈ *X* and λ ∈ (0,1), we have λ***x*** + (1 − λ)***y*** ∈ *X*. A set in ℝ^*n*^ is a *convex polyhedron* if it is an intersection of finitely many closed half-spaces in ℝ^*n*^. A *cone* is a set *X* ⊂ ℝ^*n*^ with the property that ***x*** ∈ *X* implies λ***x*** ∈ *X* for all λ ∈ ℝ and the cone is *pointed* if λ***x*** ∈ *X* if and only if λ ≥ 0. A *convex polytope* is a bounded convex polyhedron.

A vertex of a convex polyhedron 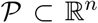 is a point 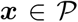 such that there exists a vector c with the property that ***c · x*** > ***c · y*** for all 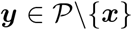. The space of feasible solutions of a Linear Program form a convex polyhedron. A standard result from Linear Programming states that the vertices of this polyhedron coincide exactly with the basic feasible solutions of the LP. Lastly, for a cone defined by

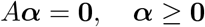

we define its support by

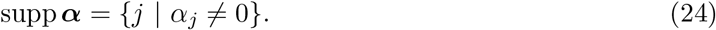

A standard result from LP is that the spanning rays of such a cone are those vectors in the cone with minimal support.

### 2.5 Definition of Elementary Growth States (EGSs)

The balanced growth equations (23a) may be succinctly written as

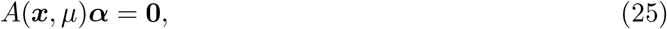

where

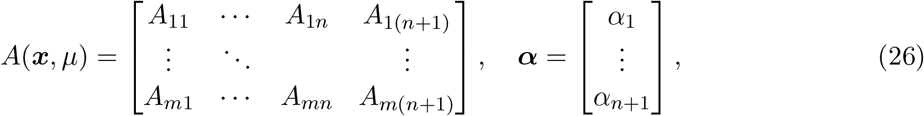

and for *k* ∈ {1,…,*m*} and *j* ∈ {1,…,*n*} we have

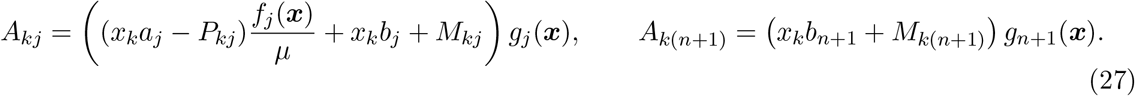

This is a system of linear equations in ***α***. Because the equation (23b) is also linear in *α*_*n*+1_, we can incorporate it in the system by introducing a new matrix,

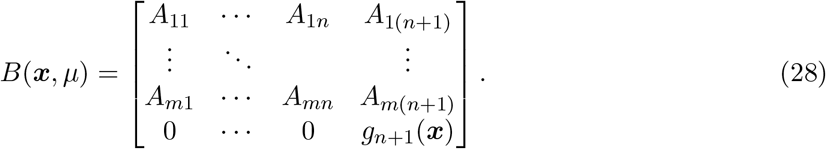

With these definitions, the balanced growth equations (23a)–(23d) read

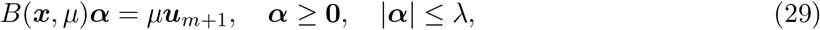

where ***u***_*m*+1_ is the unit vector: [0,…,0,1]^*T*^.

#### Remark 1.

*The rank of the matrix B*(***x***,*μ*) *may in principle depend on the choice of **x** and μ. Certainly, rows corresponding to a metabolite with zero concentration, x_k_* = 0, *are no valid constraints. If these rows are left out, it is clear, considering the definition of the elements of B*(***x***,*μ*) *in (27), that the set of ***x*** and μ for which B*(***x***,*μ*) *does not have full rank has measure zero—even if the original stoichiometry matrices may have lower rank. We therefore from now on assume, that the rank does not depend on the choice of **x** and μ. Most arguments that follow are for fixed **x**, and so we will require that the rank does not depend on μ*.

To introduce Elementary Growth States, we fix ***x*** and *μ*, and ignore the ribosome allocation inequality |***α***| ≤ λ for now. Denote the row vectors of *B* = *B*(***x***, *μ*) by 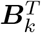, *k* = 1,…,*m* + 1. The set formed by all positive solutions to the first *m* equations,

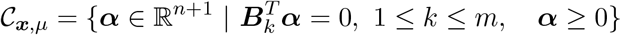

is a pointed polyhedral cone. The cone becomes a convex polytope if we restrict to those vectors ***α*** in 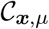 such that *α*_*n*+1_*g*_*n*+1_(***x***) = *μ*. Figure 1**a** gives the reader a sketch of this construction. Let us define

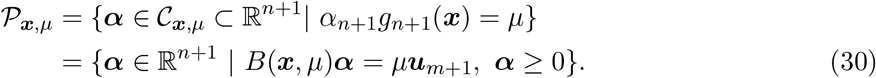

**Figure 1:**
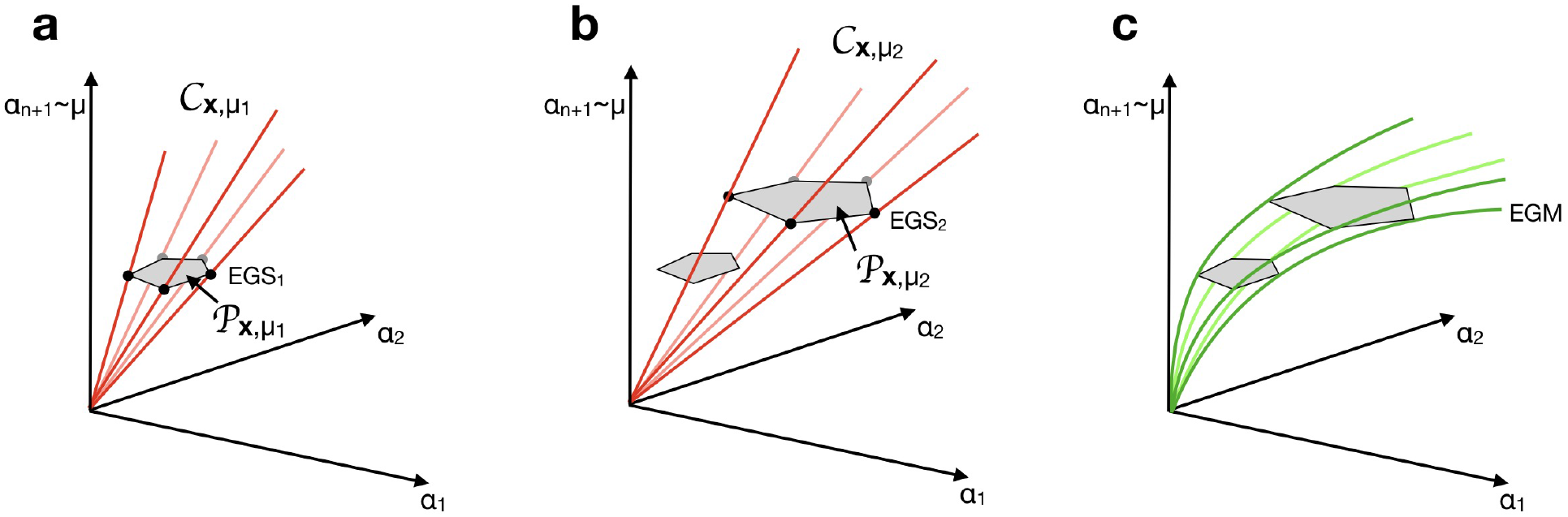
Construction of Elementary Growth States and Elementary Growth Modes. a) For fixed metabolite concentrations ***x*** and growth rate *μ*_1_, the balanced growth equations (23a) and (23c) define a cone 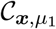. Intersecting this cone with (23b) defines a polytope 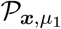 containing all balanced growth solutions. Its vertices are the EGSs at this growth rate and metabolite concentrations. b) Changing the growth rate from *μ*_1_ to *μ*_2_ changes the cone to 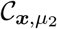 and the polytope to 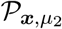, because (23a) depends on *μ*. (A similar change would be seen if ***x*** had been changed.) c) The EGSs change continuously with *μ* (and also with ***x***, not shown), but their support does not. The green lines, connecting the vertices in the different polytopes, together form the EGMs.

#### Definition 1.

*For a given set of metabolite concentrations **x** and growth rate μ, let* 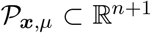 *be the corresponding convex polytope. Then the* Elementary Growth States with metabolite concentrations ***x*** and growth rate *μ are the vertices of this polytope*.

Each EGS ***α*** ∈ ℝ^*n*+1^ is thus a vertex of 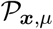 and therefore a basic feasible solution of the Linear Program (29). It has a corresponding feasible basis *D* such that *B_D_*(***x***,*μ*)***α***_*D*_ = *μ**u***_*m*+1_ and *α_j_* = 0 for all *j* ∉ *D*. Moreover, for each *D, B*_*D*_(***x***, *μ*) is square (after restricting to a set of linearly independent rows) and nonsingular. The support supp ***α*** for an EGS satisfies supp ***α*** ⊆ *D*.

EGSs are quintessentially objects that depend on the chosen metabolite concentration vector ***x*** and growth rate *μ*. For each choice, the polytope 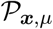 will be different, with different vertices. This is because the constraint equations that define this polytope include the stoichiometry, the kinetics of enzymatic reactions, and the metabolite concentrations of the cell.

With the definition of EGSs, we immediately have the following fundamental result.

#### Theorem 2.

*Any balanced growth solution with growth rate μ is a convex combination of EGSs with the same growth rate and the same metabolite concentrations*.

*Proof*. For fixed ***x*** and growth rate *μ*, any balanced growth solution satisfies *B*(***x***,*μ*)***α*** = *μ**u***_*m*+1_ and ***α*** ≥ 0. The vectors that satisfy these relations form precisely the polytope 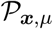. This polytope is the convex hull of its vertices, the EGSs.

A convex combination of EGSs with different growth rates does not automatically satisfy the balanced growth equations, i.e., if ***α***_1_, ***α***_2_ are two EGSs with growth rates *μ*_1_ and *μ*_2_, then the convex combination *t**α***_1_ + (1 − *t*)***α***_2_ does not necessarily satisfy

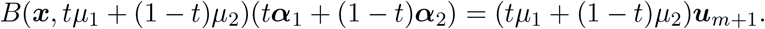

The main reason is that *B*(***x***,*μ*) changes with *μ*. For fixed ***x***, each *μ* defines a different polytope 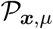, see Figure 1**b**

We may assume without loss of generality that *B*(***x***,*μ*) has maximal rank *m* + 1. If not, we can, without consequence, delete rows from *B* that are linearly dependent. This would mean that in balanced growth, not all metabolic concentrations are independent: there are ‘moieties’, but not exactly in the classical stoichiometric fashion. There is always dilution by growth. When selecting a feasible basis *D*, the last column in *B* should always be kept, since it enforces *α*_*n*+1_*g*_*n*+1_(***x***) = *μ*, and ensures that the ribosome is made; without ribosomes, there are no enzymes.

#### Theorem 3.

*If a vector **α** is an EGS, then its support supp **α** corresponds to a subnetwork that has a number of metabolic reactions that is less or equal to the number of independent metabolites. Each set D that is a feasible basis for an EGS, and for which D is also its support, has exactly as many metabolic reactions as independent metabolites*.

*Proof*. We fix ***x*** and *μ* throughout, and thereby also the polytope 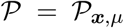. The EGSs are the vertices of 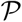, and equivalently the basic feasible solutions of the LP. (A standard LP has an objective function, but that does not concern us here. For now, we only need to focus on the basic feasible solutions.)

Each basic feasible solution has a corresponding feasible basis *D*, so that *B_D_*(***x***,*μ*) is square and invertible. Note that |*D*| must therefore be equal to the number of independent rows of *B*. Independent rows are given by the Balanced Growth equations of independent metabolites with non-zero concentrations. The vector 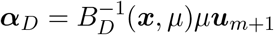 has length |*D*|. For all *j* ∉ *D* we have *α_j_* = 0, but even for *j* ∈ *D* we can have *α_j_* = 0, so that we have at most *D* non-zero *α_j_*’s. Certainly *n* + 1 ∈ *D*, because it corresponds to the ribosome, without which enzymes cannot be synthesised. That leaves |*D*| − 1 elements *j* ∈ *D*\{*n* + 1}, each of which corresponds to a unique *α_j_*, and thus to a unique enzyme. The number of active reactions, corresponding to non-zero *α_j_*’s, is thus less or equal to the number of independent metabolites (in the sense described above) with non-zero concentrations.

If *D* is also the support of *α*, then the number of non-zero *α_j_*’s is equal to |*D*|, so that the number of active reactions is exactly equal to the number of independent metabolites with nonzero concentration.

#### Biological interpretation and consequences

Not all protein expression profiles lead to balanced growth: at least the essential enzymes needed for growth on a set of nutrients need to be made. There exist also protein profiles in which the same metabolites are being produced in multiple different ways, or in which different modes of metabolism are active at the same time, such as with overflow metabolism. Such networks show redundancy.

We have identified Elementary Growth States as the simplest, non-decomposable, cellular states that allow balanced growth. Any balanced growth state may be decomposed into EGSs that share the same growth rate and metabolite concentrations, but have different ribosomal allocations and use a different set of reactions. EGSs, as the simplest cellular phenotypes that include the concentrations and synthesis rates of all cellular components, are therefore the fundamental building blocks of balanced cellular growth.

Note that this set of building blocks is not yet invariant. For example, changes in the growth rate, the metabolite concentrations, or the kinetic parameters will change the EGSs. Since a set of invariant building blocks has proven very useful in FBA-theory (EFMs), we will now define Elementary Growth Modes that will indeed be invariant.

### 2.6 Elementary Growth Modes are equivalence classes of Elementary Growth States

The support of an EGS, defined in (24), induces a natural equivalence class of EGSs which all share the same support and hence constitute steady state networks with the same enzymes. This equivalence class does *not* depend on the *μ* and ***x*** for which the individual EGSs were calculated.

#### Definition 2.

*Two EGSs, **α***_1_(***x***_1_,*μ*_1_) *and* ***α***_2_(***x***_2_,*μ*_2_), *are said to be equivalent if they have the same support. The equivalence class to which **α***_1_ *belongs is denoted by* [***α***_1_(***x***,*μ*)], *and is called an Elementary Growth Mode*.

The set of EGMs,

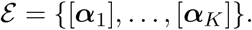

contains all essentially different minimal networks that can sustain balanced growth, each with its unique set of participating reactions. In mathematical terms, the set of all EGMs, 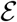, is the quotient set of all EGSs under identification of EGSs with the same support.

Note that for each ***x*** and *μ* individually, a representative EGS for each EGM equivalence class can be calculated using standard algorithms. It is just a matter of finding the vertices of the polytope 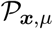 defined in (30), a task for which computational algorithms are already available. These representatives will change with ***x*** and *μ*, but their support (and thus the EGM that they belong to) will not. It is however possible that an EGM has a feasible representative at some set of metabolite concentrations and growth rate, but not for another set.

We here prove that an EGS can be continuously extended both in an open neighbourhood around *μ* and in a neighbourhood around ***x***. This shows that if the whole-cell model allows a steady state at some fixed ***x*** and *μ*, then steady states also exist with the same participating reactions for metabolite concentrations and growth rates close to the fixed values, see Figure 1**c**.

In other words, given a representative of an EGM, this EGM also has a feasible representative in a small neighbourhood.

#### Theorem 4.

*For a given growth rate μ*_0_ *and set of metabolite concentrations **x***_0_, *there exists an open neighbourhood U such that for all* (***x***,*μ*) ∈ *U, each EGS with support equal to its feasible basis, **α***(***x***_0_,*μ*_0_), *can be continuously extended to a vector **α***(***x***,*μ*) *that solves the balanced growth equations and belongs to the same EGM*.

*Proof*. Choose an arbitrary EGS: ***α***_0_ ∈ ℝ^*n*+1^. We know that it solves

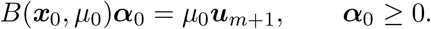

The EGS has a feasible basis *D*, and without loss of generality we restrict *B*(***x***_0_, *μ*_0_) to the columns indexed in *D*. As discussed before, we can select a set of rows corresponding to independent metabolites with non-zero concentrations such that the resulting matrix is square and invertible. We may therefore choose *n = m* and the dimension of *B* to be (*m* + 1) × (*m* + 1).

We would like to apply the Implicit Function Theorem to see that there are continuously differentiable functions 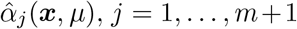, such that in an open environment of (***x***_0_, *μ*_0_), the Balanced Growth Equations are still met: 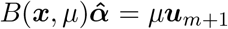. Since the 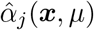 are continuous, and since no *α_j_*(***x***_0_, *μ*_0_) is equal to zero by the assumption in the Theorem, we can then also choose a neighbourhood in which 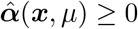. This 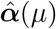 would thus indeed be a continuous extension of the EGS that belongs to the same EGM, since its support does not change.

Let *s* be the number of components in ***x***. For the Implicit Function Theorem we need a function *F*: ℝ^*m*+1^ × ℝ^*s*+1^ → ℝ^*m*+1^ that is zero at (***α***_0_, ***x***_0_, *μ*_0_). For this function, we can use the Balanced Growth equations as components: the first m are given by

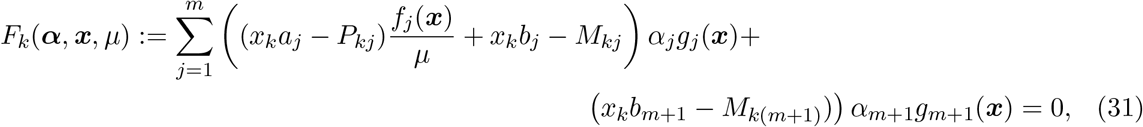

where *k* = 1,…,*m*. The last component is given by

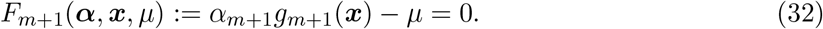

Let us check the conditions for the Implicit Function Theorem:

- The function *F*: ℝ^*m*+1^ × ℝ^*s*+1^ → ℝ^*m*+1^ with components given by (31) and (32) is continuously differentiable in ***α, x***, and *μ* in a neighbourhood of (***α***_0_, ***x***_0_, *μ*_0_).
- By assumption, (***α***_0_, ***x***_0_, *μ*_0_) is a solution of the balanced growth equations: *F*(***α***_0_, ***x***_0_, *μ*_0_) = 0.
- The entries of the Jacobian of *F* at (***α***_0_, ***x***_0_, *μ*_0_) with respect to ***α*** are, for *k* = 1…,*m*, given by

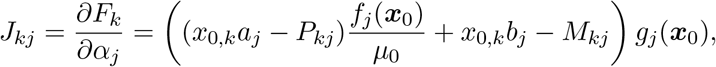

and the last row of the Jacobian is zero except for the last entry, which is *g*_*m*+1_(***x***). Note that this Jacobian is exactly the matrix *B*(***x***_0_,*μ*_0_), which was invertible by construction.

The IFT may therefore be invoked, which shows that for each EGS at (*x*_0_,*μ*_0_), with support equal to its feasible basis, there is an open neighbourhood around (***x***_0_,*μ*_0_) such that the EGS can be continuously extended. By taking the intersection of these neighbourhoods we can indeed find a neighbourhood in which all of the EGSs can be extended.

The previous theorem shows that the set of EGMs is largely conserved when the growth rate or the metabolite concentrations are changed. Indeed, the functions 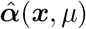 that were found in the previous proof in principle give rise to balanced growth vectors at a wide range of growth rates *μ* and concentrations ***x***. However, these vectors are not necessarily all EGSs, because the positivity constraint ***α*** ≥ 0 may be broken. If one of the components 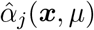 becomes smaller than zero, then the corresponding EGM ceases to exist. We will use this fact in Theorem 5.

#### Biological interpretation and consequences

If a cell is able to sustain itself at particular fixed metabolite concentrations and at a certain growth rate, then it is still able to maintain itself at slightly perturbed concentrations or a different growth rate, using the same enzymes and hence the same metabolic network. This network structure is defined to be the Elementary Growth Mode. These EGMs therefore form the set of invariant building blocks that we were after, analogous to the EFMs in FBA-models.

Indeed, the EGM may be used at different metabolite concentrations and balanced growth rates as long as none of the constraints is violated. These constraints will eventually be violated however: in particular, a higher growth rate may either require a ribosomal fraction to become negative (which is physically impossible), or it may become too large, and violate |***α***| ≤ λ.

### 2.7 Maximal growth rate is attained in EGMs

We now prove that maximal growth rate is attained in an EGS, and hence in an EGM. Until now we have ignored the upper bound on the ribosome allocation, expressed in

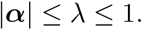

From now on, we will consider this bound, and study the optimisation problem of finding the maximal balanced growth rate for the whole-cell model (16), phrased here as

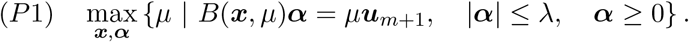

Since *μ* is part of the construction of the solution polytope 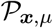, we formulate a different optimisation problem, in which the metabolite concentrations are fixed, and we do not maximise the growth rate but we minimise the total fraction of the ribosome necessary to attain a given growth rate,

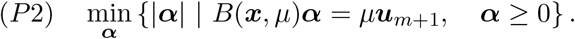

Problem (P2) should be considered for growth rates *μ* that are relevant for (P1); for instance, for the growth rate *μ*_max_ that maximises (P1). Also note that (P2) may be reformulated as

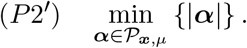

Figure 2 gives some intuition behind the proof of the following theorem.

**Figure 2:**
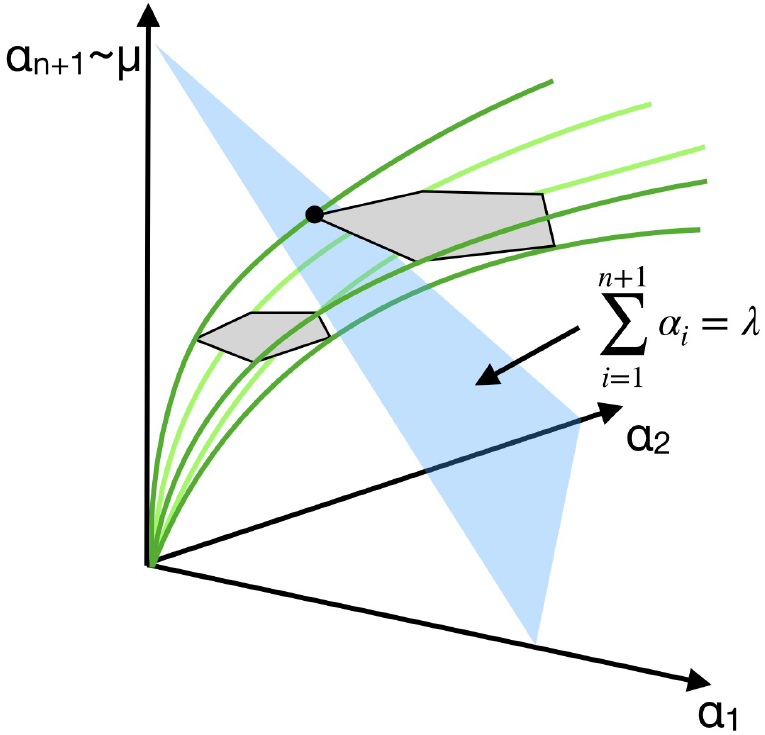
Intuition behind the proof of Theorem 5. As *μ* is increased, the polytope 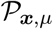 cuts through the ribosome allocation constraint 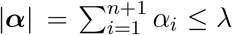, with |***α***| = λ shown in blue. Exactly at the maximal growth rate, the intersection of the polytope with |***α***| = λ is reduced to one vertex (black dot), i.e., one EGS.

#### Theorem 5.

*For each choice of* λ, (*P1*) *is maximised in an Elementary Growth State*.

*Proof*. Let us consider the maximisation problem for an arbitrary fixed set of metabolite concentrations ***x*** (inspired by the proofs from (Wortel et al., 2014; de Groot et al., 2019)). For this fixed ***x*** we will prove that the problem is always maximised in an EGS. This is enough to prove the theorem because this means that the growth rate is also maximised in an EGS if we would have picked the optimal metabolite concentrations ***x***.

We only need to consider those ***x*** for which (P1) has a nonempty set over which to maximise. The guiding insight for this proof is that, for each fixed value of *μ* such that the polytope 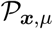 is nonempty, the minimum of (P2) is attained in a vertex of this polytope, i.e., in an EGS with growth rate *μ*. (This is true for any linear objective function of ***α***).

Let *μ*_max_ be the value of the growth rate that maximises (P1), with maximiser ***α***^max^, and |***α***^max^| =: λ^max^ ≤ λ. It is clear that 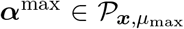. For *μ* > *μ*_max_, 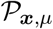 is either empty, or it is not, so we distinguish two cases.

Case 1: If 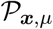 is not empty for some *μ* > *μ*_max_, then |***α***| > λ for all 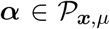; if not, those vectors would satisfy all the constraints of (P1), so *μ*_max_ could not have been maximal. Moreover, vertices of 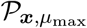 change continuously as a function of *μ* in a neighbourhood of *μ*_max_ (Theorem 4), and since ***α***^max^ is a convex combination of those vertices, ***α***^max^ itself also changes continuously. Finally, because |***α***| is a continuous function of ***α***, the change in |***α***^max^| will also be continuous. Adding this to the observation that |***α***| > λ for all 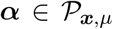, *μ* > *μ*^max^, we can conclude that |***α***| ≥ λ for all 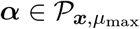.

Since also |***α***^max^| ≤ λ, we must have |***α***^max^| = λ, and ***α***^max^ is thus a minimiser of (P2) for *μ* = *μ*_max_. The minimum of (P2) is attained in a vertex of the polytope, so we can choose ***α***^max^ such that it is an EGS.

Case 2: If 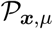 is empty for all *μ* > *μ*_max_, then all ***α*** that solve *B*(***x***, *μ*)***α*** = *μ**u***_*m*+1_ for *μ* > *μ*_max_, must have at least one index *j* such that *α_j_* < 0. However, we also know that the polytope was nonempty for *μ* = *μ*_max_, and a nonempty polytope must have at least one vertex. We can thus choose ***α***^max^ to be such a vertex, and the maximal growth rate is thus indeed maximised in an EGS.

Note that in both cases there could, in principle, be more than one choice for ***α***^max^: in Case 1 there could be an entire edge of the polytope that minimises (*P*2) for *μ* = *μ*_max_, and in Case 2 the polytope at *μ* = *μ*_max_ could have more than one vertex. This however does not contradict the theorem since the theorem does not state that (*P*1) is *only* maximised in an EGS, but rather that it *is* maximised in some EGS. This is of course a standard situation in Linear Programming theory.

The proof shows that there are two situations in which the growth rate is maximised. Either the constraint |***α***| ≤ λ is hit, or one of the *α_j_* = 0 and becomes negative at larger growth rates, causing the network to lose an enzymatic reaction and thereby ‘disintegrate’ (it cannot support balanced growth anymore).

We noted that the maximiser of (*P*1) does not have to be unique. It is however very unlikely to be non-unique if enough information about the enzyme saturation functions, *f_j_*(***x***), and the volumetric parameters, *ρ_k_, σ_j_*, is provided. Having a non-unique maximiser means that there are at least two EGMs, ***α***_1_, ***α***_2_ that have exactly the same maximal growth rate, thus these EGMs either both have |***α***_1_| = |***α***_2_| = λ or both become infeasible, at *μ* = *μ*_max_. However, the two EGMs must by definition have a different support, such that there is at least one reaction *j* that is only used in the first EGM, and a reaction *j*′ that is only used in the second. The Balanced Growth equations for an EGM, and thus the maximal growth rate for that EGM, depend on the parameters of all participating reactions. The different parameters (e.g., different stoichiometry, volumetric constants, and enzymatic parameters) for the reactions *j* and *j*′, will thus influence the maximal growth rate, making it very unlikely that the maximal growth rates for the two EGMs are exactly the same.

We also remark that (P1) is a problem for all metabolite concentrations and all ribosome allocations, whereas (P2) is a problem only for the ribosome allocation. We can also consider an optimisation problem for the metabolite concentrations alone: if we pick an EGM we can maximise the growth rate in that EGM by varying ***x***. In this stage we do not know whether this optimum is unique, or what the convexity properties are of this problem.

The proof of Theorem 5 also shows that increasing λ must coincide with increased *maximal* growth rate.

#### Corollary 6.

*Consider (P1) for two values of* λ, 0 < λ_1_ ≤ λ_2_, *and denote* 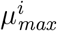 *the maximal growth rate attained with* λ = λ_*i*_, *i* = 1,2. *Then* 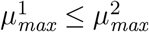.

The converse of this corollary does not necessarily hold: if *μ*_1_ ≤ *μ*_2_ and 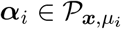 for *i* = 1, 2, it does not follow that |***α***_1_1 ≤ |***α***_2_|.

#### 2.7.1 Adding additional linear enzymatic constraints results in minimal convex combinations of EGMs

In a cell, protein concentrations are bounded. This is the direct consequence of cells having membranes with finite areas, compartments with finite volumes and proteins occupying volume and space. Accordingly, each membrane and compartment in a cell has a finite capacity to contain proteins, effectively setting constraints to the protein concentrations in cells. These protein constraints are often modelled as upper bounds to weighted sums of enzyme (and ribosome) concentrations: 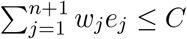. To include these bounds in the optimisation problem, we need to find expressions for the enzyme concentrations *e_i_* in terms of the *α_j_*’s.

The steady state value of the enzyme concentrations is 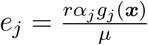 (note that in steady state, this identity also holds for the ribosome concentration, *j* = *n* + 1, so we don’t have to treat it separately). Using the steady state ribosome concentration (19), which reads

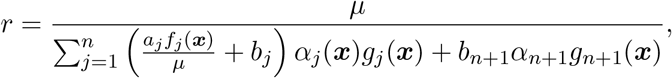

this becomes

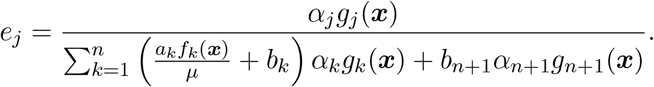

We can use this to rewrite a linear constraint on the enzyme concentrations into a linear constraint on the *α_j_*’s, by rewriting 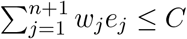 to

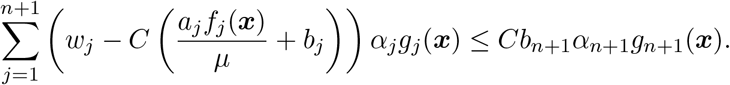

These are inequalities with the same type of dependency on ***α*** as before, and may be added to the optimisation problem. In a similar way, a linear combination of enzymes and the ribosome will produce inequalities with *α_j_* of the same type.

##### Theorem 7.

*The optimisation problem (P1), supplemented with K linear constraints*

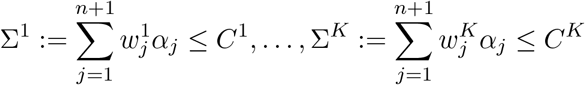

*has a solution that is a convex combination of at most K* + 1 *Elementary Growth States*.

*Proof*. In the proof of Theorem 5 we showed that the maximal growth rate *μ* will be attained by a vertex of the polytope 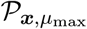. Here we show that this proof can still be used for a slightly different polytope, defined by

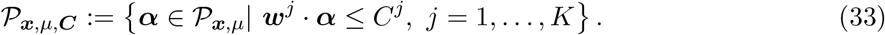

Following the proof of Theorem 5 we can separate two cases: in Case 1 there is a *μ* > *μ*_max_ for which the polytope is nonempty, and in Case 2 the polytope is empty for all *μ* > *μ*_max_. The two cases can be handled exactly as in the proof of Theorem 5 so that we know that the maximal growth rate under the additional constraints will be attained in a vertex of 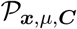. Therefore, if we can describe the vertices of the new polytope, we are done.

Vertices are 0-dimensional faces of the polytope. In an *n* + 1-dimensional space, the vertices should therefore, just as before, satisfy *n* + 1 equalities. Of these equalities, *K* could come from the newly added constraints. That means that a vertex of the new polytope should satisfy at least *n* + 1 − *K* equalities in the old polytope 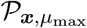. In other words, the vertex was part of a *K*-dimensional face of the old polytope and therefore it is a convex combination of at most *K* + 1 of the 0-dimensional EGSs.

##### Biological interpretation and consequences

EGMs are the self-fabricating networks that maximise balanced growth rate. By minimising the number of enzymatic reactions, the investment in each of the remaining reactions may be maximised, leading to the highest possible overall synthesis rates, and thus the maximal growth rate. While maximising the growth rate, biophysical constraints may be hit. Such constraints may often be formulated as linear combinations of enzyme concentrations, such as the total enzyme concentration in the cytosol, total enzyme concentration in the membrane, and so on. When constraints are hit, the maximiser is not necessarily an EGM, but possibly becomes a convex combination of EGMs. The number of such EGMs found in the optimum is less or equal to the number of constraints. Constant osmotic pressure, assumed in (15), is the first constraint. Since the number of possible physical constraints is limited, the theory predicts that optimal steady state behaviour is simple, with a small number of EGMs active at any time.

### 2.8 The relation between EFMs and EGMs

Elementary Flux Modes (EFMs) are the fundamental building blocks of genome-scale stoichiometric models (Schuster et al., 2000). EFMs are minimal in the sense that all the reactions in an EFM are essential for a steady state flux. The EFMs can be found as the extreme rays of the flux cone (see Figure 3), if reversible reactions are split in two separate irreversible reactions. The set of all feasible flux vectors, given the reaction stoichiometry, can then be described as the set of all convex combinations of EFMs.

**Figure 3:**
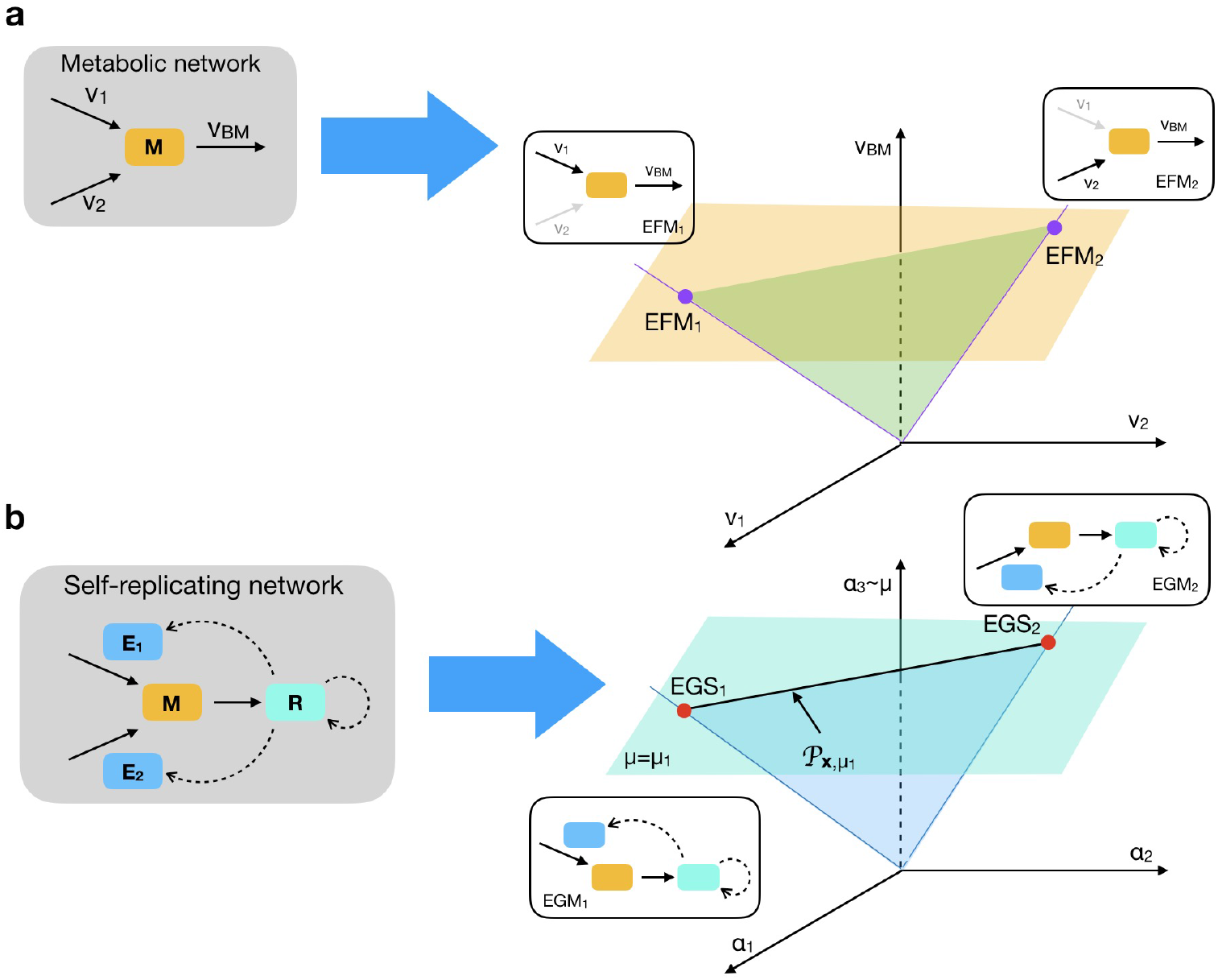
Comparison between EFMs and EGSs/EGMs for a simple metabolic network (a) with one metabolite M, and three reactions, and the corresponding self-replicating network (b) with two enzymes E_1_ and E_2_ and the ribosome R. In case (a) the cone of possible steady state flux solutions is a plane through the origin, bounded by *v*_1_ ≥ 0 and *v*_2_ ≥ 0. Restricting to solutions with those with a prescribed biomass flux vbm (orange horizontal plane) gives a line segment bordered by two EFM vertices. In EFM_1_, reaction *v*_1_ participates; in EFM_2_, reaction *v*_2_. In (b) for given metabolite concentration *x* and growth rate *μ*_1_, the balanced growth solutions for the corresponding self-replicating network also form a line segment of a cone, again spanned by two vertices, the EGSs at these fixed values of *x* and *μ*. Only this line segment contains balanced growth solutions, the rest of the cone does not. In EGS_1_ only enzyme 1 and the ribosome are expressed, while in EGS_2_, enzyme 2 and the ribosome are expressed.

To relate EFM theory to EGM theory, we start with stoichiometric matrices *P* and *M* of the whole-cell model (16). Choosing an EGS requires first choosing metabolite concentrations ***x*** and a growth rate *μ*, and then choosing a feasible basis *D* which gives rise to a square invertible matrix *B_D_*, corresponding to a vertex of the polytope 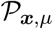. We may therefore also restrict the stoichiometry matrices to *P_D_* and *M_D_*, accordingly.

In genome-scale metabolic networks that only model the metabolic part and not the enzymatic and ribosomal part of self-fabrication, a virtual biomass reaction is added. This reaction consumes all components necessary for cell synthesis in (approximately) the right proportions. The growth rate is assumed to be proportional to the rate of the biomass reaction. In the following theorem we find, for each EGS, a suitable biomass reaction, such that the metabolic part of the network (*P_D_*) appended with this virtual reaction has one EFM. The flux values of this EFM approximate the flux values of the EGS. The correction is of order *μ*, which is usually several orders of magnitude smaller than the flux values.

#### Theorem 8.

*Let **α** be an EGS for metabolite concentrations **x** and growth rate μ, with feasible basis D. Let v* ∈ ℝ^*n*^ *be the corresponding flux vector and **w*** ∈ ℝ^*n*+1^ *the corresponding enzyme/ribosome synthesis flux vector. Let ϕ* = *M***w**, where M is the enzyme synthesis stoichiometry matrix, and assume that [*P_D_* − *ϕ*] *has full rank. Then we may write*

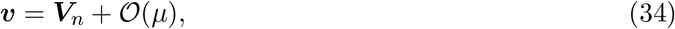

*where* 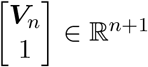 *is the unique appropriately scaled EFM of the stoichiometric matrix* [*P_D_* − *ϕ*].

*Proof*. Let *P* = *P_D_*. At balanced growth, the flux vectors ***v*** and ***w*** satisfy

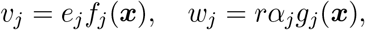

where

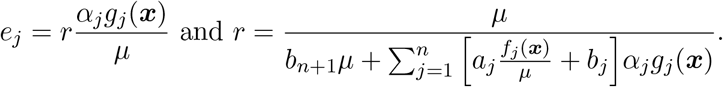

These vectors satisfy the steady state equations *ẋ* = 0, i.e.,

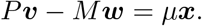

Setting *ϕ* = *M****w*** and *P_ϕ_* = [*P* − *ϕ*], we can see that ***v*** is given by the linear set of equations,

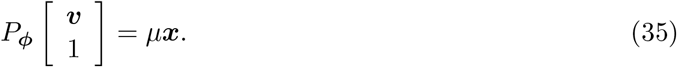

We analyse this problem for fixed ***x*** and *ϕ*, which must have solutions, since ***v*** is already one such solution. We call ***u*** ∈ ℝ^*n*+1^ a solution to *P_ϕ_****u*** = *μ****x***. Since we have assumed that *P_ϕ_* has full rank, we know that there must be a one-dimensional solution space. We further exploit that *P_ϕ_* has full rank by using row-reduction to write 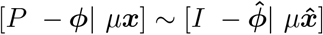. Row-reduction does not change the solution space, so, if we let ***u***_*n*_ denote the first *n* entries of ***u***, 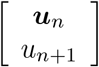 solves 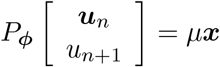 if and only if it also solves 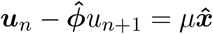. The latter is solved by 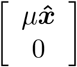, so the first must be too, such that this gives us a particular solution. The one-dimensional solution space is then found by adding a scalar multiple of a vector, *u*\*, from the nullspace of *P_ϕ_*. So,

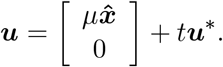

The vector ***u***\* is the EFM of the matrix *P_ϕ_*. Let *t*\* be chosen such that 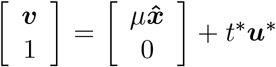.

(This is always possible, since we already have a solution ***v***.) Setting 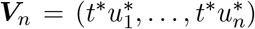, we deduce

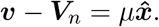

The theorem states that [*P_D_* − *M****w***] should have maximal rank. This requires at least that rank *P_D_* ≥ *m* − 1. This does not follow directly from the fact that *B_D_*(***x***, *μ*) has full rank; one can easily make a counterexample. We do expect that *P_D_* has full rank in many cases.

#### 2.8.1 The approximating EFM is constant if the precursor consumption rates change proportionally

The relevant EFM for an EGS is thus the vector ***V*** spanning the null space of the matrix *P_ϕ_*, excluding the last element *V*_*n*+1_. The total precursor consumption rate, *ϕ* = *M****w***, is part of the EGS solution and is therefore not known a priori. In this section, however, we assume that the relative precursor consumption rates are constant, making *ϕ* a fixed vector, up to a proportionality constant. We show that under this assumption, the EGS is approximated by the same EFM for all metabolite concentrations and growth rates.

##### Theorem 9.

*Let* ***α***^0^ *be an EGS for metabolite concentrations **x***^0^ *and growth rate μ*^0^, *with feasible basis D. Let **v***^0^ ∈ ℝ^*n*^ *be the corresponding flux vector and **w***^0^ ∈ ℝ^*n*+1^ *the corresponding enzyme synthesis flux vector. Let ϕ*^0^ = *M****w***, *where M is the enzyme synthesis stoichiometry matrix, and assume that* [*P_D_* − *ϕ*^0^] *has full rank. Let **V**_n_ be the EFM that approximates the flux values of **α***^0^, *according to Theorem 8. Moreover, assume that ϕ*(***x***,*μ*) = *h*(***x***,*μ*)*ϕ*^0^ *with h some scalar function. Then, for all μ and **x** such that D is a feasible basis, the flux values **v***(***x***,*μ*) *are approximated by the same EFM*:

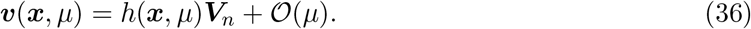

*Proof*. Following the proof of Theorem 8 we know that the flux values of an EGS are approximated by the first *n* coordinates of a vector that is in the nullspace of *P_ϕ_* = [*P* − *ϕ*]. Since 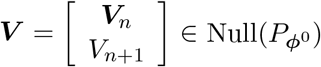, we can set ***V***_*ϕ*(***x***,*μ*)_:= (*h*(***x***, *μ*)*V*_1_,…,*h*(***x***, *μ*)*V_n_,V*_*n*+1_)^*T*^. We have

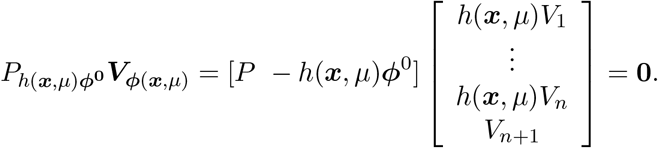

Note that the first *n* entries of ***V*** are all multiplied by the same scalar. So, the ratios of all metabolic reaction rates (not including the virtual biomass reaction) are constant.

In Figure 4 we provide evidence that the amino acid composition of cells growing in different media are practically equal. With the above theorem, this indicates that flux values of EFMs likely closely resemble those of EGSs, provided that these EFMs were calculated for an accurately estimated biomass reaction.

**Figure 4:**
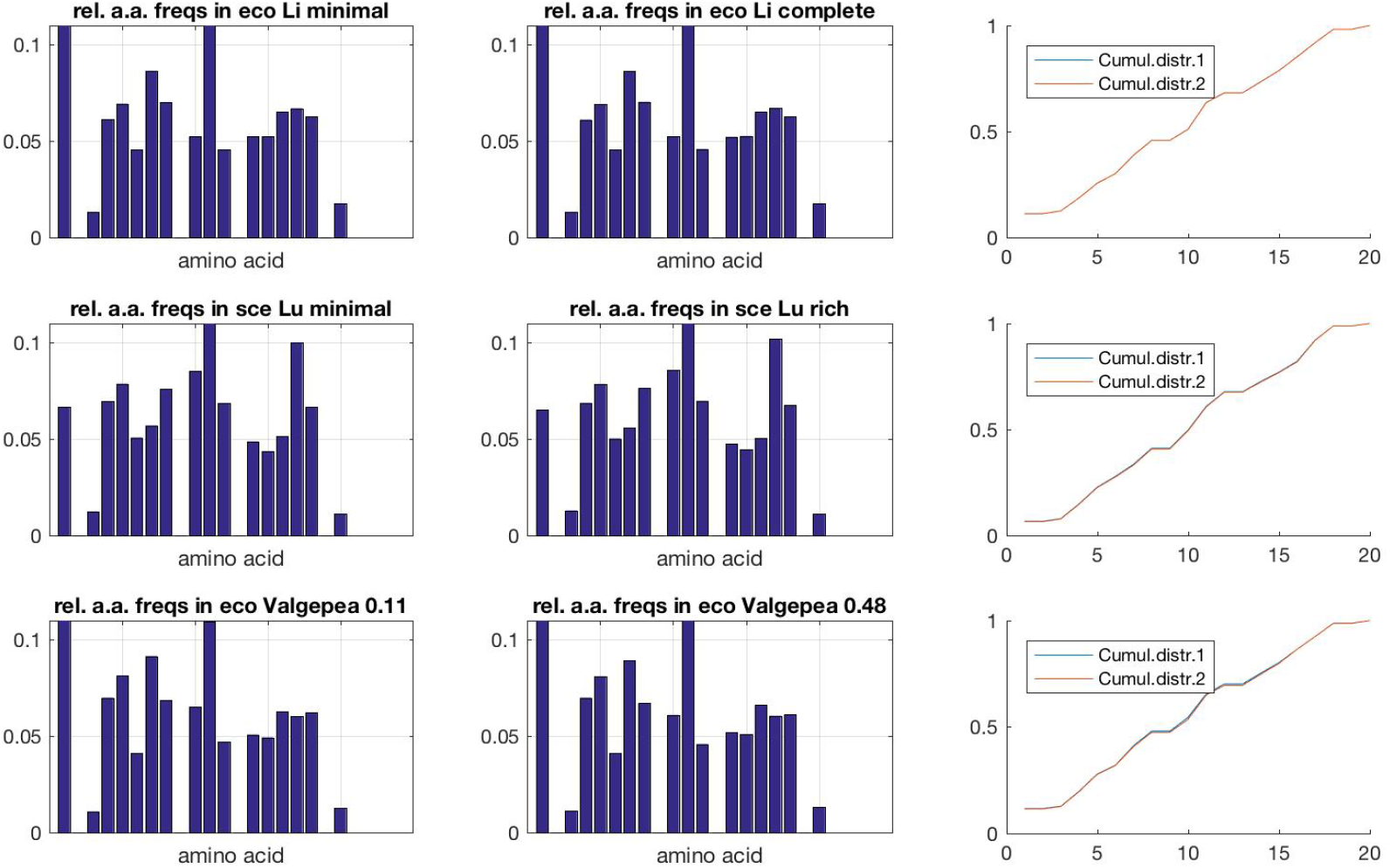
Side-by-side comparisons of relative amino acid frequencies in growing cells, in minimal and rich media. Bar plots give relative amino acid frequencies in minimal (left column) and rich medium (middle column). Right column illustrates the cumulative frequency distributions used in the Kolmogorov-Smirnov (KS) test. KS statistic values (which measure the maximal difference between these two cumulative distributions) for the three studies are 3.5 · 10^−4^, 3.6 · 10^−3^ and 9.3 · 10^−3^, indicating that these distributions can hardly be distinguished (as is evident from the plots). Data (available at `proteomaps.net`) from (Li et al., 2014) (top row; *E. coli*) (Lu et al., 2007) (middle row; *S. cerevisae*) and (Valgepea et al., 2013) (bottom row; *E. coli*). See (Teusink et al., 2006) for another experimental example from *Lactobacillus plantarum*.

#### 2.8.2 The dependence of EGSs on growth rate

The constant approximation of EGS flux values up to a correction of order *μ* by an EFM can be used to investigate the dependence of an EGS, ***α***, on the growth rate. If we make the additional assumption that the dilution of metabolites is negligible, we can even make the *μ*-dependence explicit.

##### Theorem 10.

*Let* ***α***^0^ *be an EGS for metabolite concentrations **x***^0^ *and growth rate μ*^0^, *with feasible basis D. Let* 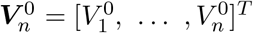 *be the EFM that approximates the flux values of **α***^0^, *according to Theorem 8. Moreover, assume that ϕ*(***x***,*μ*) = *h*(***x***,*μ*)*ϕ*^0^ *with h some scalar function, and that the dilution of metabolites is negligible compared to metabolic fluxes, μx_k_* = 0. *Then, there is an upper bound for the growth rate attained in this subnetwork, μ_ub_, and the dependence of the EGS **α** on μ* ∈ [0, *μ_ub_*] *is given by*

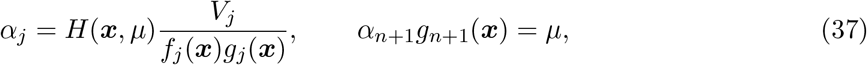

*where*

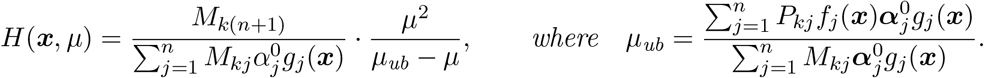

*Proof*. According to Theorem 9, the reaction rates are given by ***v***(***x***, *μ*) = (*h*(***x***, *μ*)*V*_1_,…,*h*(***x***, *μ*)*V_n_*)^*T*^. Consequently, the enzyme concentrations can be calculated by 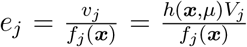, and also by 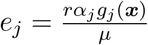. If we isolate *α_j_* from these two expressions for the enzyme concentrations, we get:

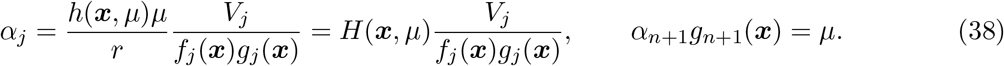

Note that the *μ*-dependent part of this expression is equal for all metabolic reactions *j* ≤ *n*. This shows that the first *n* coordinates of the EGSs change proportionally when the growth rate changes at fixed metabolite concentrations.

Given this information we can make the *μ*-dependence even more explicit, by reconsidering the balanced growth equations given in (26) and (27). Let us start from an EGS ***α***^0^(***x***_0_, *μ*_0_). We know that the EGS satisfies *A*(***x***_0_, *μ*_0_)***α***^0^(***x***_0_, *μ*_0_) = 0. For general *μ* these equations, under the assumption that the dilution of metabolites is negligible, are given by

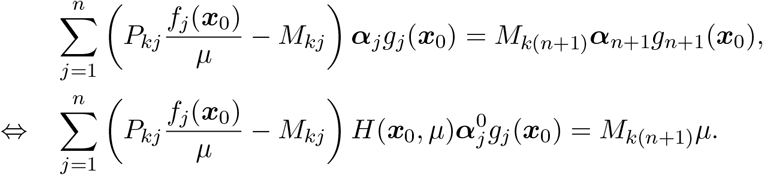

A metabolite for which *M*_*k*(*n*+1)_ = 0 yields no information on *H*(***x***_0_, *μ*), but we know that there is at least one *k* such that *M*_*k*(*n*+1)_ > 0, because the synthesis of ribosomes needs at least one building block. We pick such a *k* to isolate *H*(***x***_0_, *μ*):

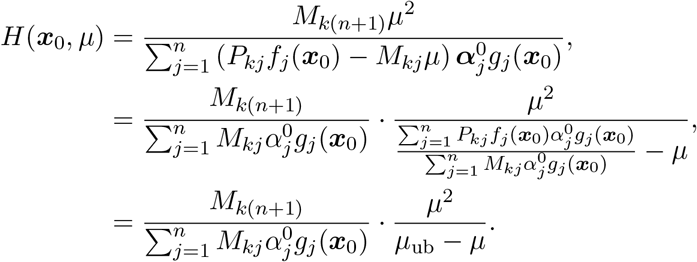

Note that *H*(***x***_0_, *μ*) is an increasing function of the growth rate, with a vertical asymptote at *μ* = *μ*_ub_. We thus see that under the assumptions that metabolite dilution is negligible and that the biomass composition remains relatively constant, the specific index set *D* gives rise to a valid EGS for the range [0, *μ*_ub_]. However, the ribosome allocation constraint |***α***| ≤ 1 will already constrain growth before *μ*_ub_ is approached. For each EGS we thus get an upper bound on, but not an accurate estimation of, the maximum achievable growth rate.

This upper bound is due to the following mechanism: the expression of an enzyme at a higher concentration leads both to a higher demand for its building blocks (because more enzyme is diluted per unit time), and to a higher supply of these building blocks (since the enzymes catalyse the production of building blocks). However, the demand for building block *k*, 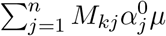, scales with *μ*, while the supply, 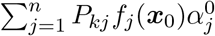, does not. The upper bound, *μ*_ub_, on the growth rate is given by that growth rate at which the expression of more protein increases the demand for building blocks more than that it increases the production of metabolites: 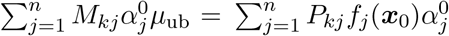. The building blocks can thus no longer be kept in steady-state.

##### Biological interpretation and consequences

The results in this section have several important implications. First of all, we can finally understand more deeply why Flux Balance Analysis has been such a good predictor of microbial metabolism. The main reason is that, when relative amino acid usage (so the relative rates of amino acid consumption for enzyme and ribosome synthesis) is constant across conditions, much of the nonlinearity in self-fabrication disappears. A linear model that disregards enzyme synthesis, but that replaces this by an accurately estimated biomass reaction, will predict flux values that are indeed close to flux values in the corresponding nonlinear model in which enzyme and ribosome synthesis are included.

Second, these constant relative amino acid consumption rates indicate why growth regulation in microbes is regulated at the amino acid level. The cell must maintain constant amino acid consumption rates, and has immediate end-product inhibitions in place in which an overabundance of a particular amino acid causes a shutdown of the synthesis of that amino acid by allosteric inhibition.

Third, we have shown that under this ‘amino-acid-assumption’, all the enzyme allocation fractions *α*_1_,…,*α_n_* change with the same factor if the growth rate and metabolite concentrations are changed. In other words, not only the EGM is kept the same (the cell keeps using the same reactions), but the relative investments in the different enzymatic reactions also remains the same! This could explain why microbes such as *E. coli* have the alarmone ppGpp which signals the depletion of amino acids, and causes a change in allocation from enzyme synthesis (i.e., having more ribosomes) to metabolism (by creating more enzymes) (Scott et al., 2014; Bosdriesz et al., 2015). Since the *relative* enzyme synthesis rates do not change with growth rate, the cell needs only to control *α*_*n*+1_ versus the rest. This dramatically simplifies the overall optimisation problem.

Fourth, under the additional assumption that the dilution rates of metabolites are negligible compared to their metabolic production and consumption rates, we identified an upper bound on the growth rate. When this upper bound is approximated, the ribosomal allocation fractions αj increase asymptotically. Therefore, the upper bound will in reality never be reached because the ribosomal allocation constraint |***α***| ≤ 1 will be limiting first. However, the upper bound still shows that the costs of growth increase nonlinearly with the growth rate, and it shows that there is a fundamental limit to self-fabrication rates.

## 3 Discussion

### A biochemical theory of balanced, unicellular self-fabrication

Self-fabrication, self-repair and phenotypic adaption to new environments are defining characteristics of autonomously living organisms. Understanding them in terms of the underlying biochemistry is a key challenge in cell biology. In this work, we focused on the average cell in a population that is growing balanced (arguably the only well-defined state in microbiology Schaechter et al. (2006)) and in a static environment. We defined the phenotype of such an average cell as the complete set of concentrations of all biochemical components, and all rates of chemical reactions. An expression for cellular growth was derived in terms of the production rates of cellular components. We then asked which phenotypes could give rise to steady state balanced growth.

To get a mechanistic understanding of how these different phenotypes can be sustained by a cell, we have derived a theory in which the quantities that are directly controlled by gene expression – the allocation fractions of the ribosome – are the free variables (inspired by Molenaar et al. (2009)). We ignored the precise mechanism how gene expression gives rise to these allocation fractions, but this does not influence the main findings of our theory. In currently used modelling approaches, the free variables are often only indirectly controllable, e.g., the reaction rates or the enzyme concentrations. By taking this new perspective, we can get a full description of the problem that needs to be solved by cells by means of gene expression, instead of only getting a description of the possible solutions of this problem.

### EGMs: the minimal phenotypes that lead to balanced growth

To describe all balanced growth phenotypes, we identified EGMs: the minimal modes of gene expression that lead to balanced growth; all possible phenotypes are convex combinations of these minimal modes. The EGMs can thus be seen as the regulatory degrees of freedom of the cell: they comprise the full capabilities of the cell in balanced growth. In contrast, one could suggest to view the expression of single enzymes as the most important degrees of freedom, but an enzyme alone can never lead to a self-fabricating system. Even an entire pathway that carries a steady-state flux does not lead to a self-fabricating system, unless it synthesises all cellular building blocks that are needed for the enzymes catalysing this pathway. Regulation should therefore not involve making numerous decisions about independent enzyme expression levels: the decision should be which EGM or combination of EGMs to express.

In one specific environment, we can calculate all minimal growth-supporting enzyme expression states, which we called Elementary Growth States. However, when the environment changes the EGSs change with it, i.e., the specific allocation fractions of the ribosome change. The cause of this change is that the saturations of enzymes and the ribosome change, and because the dilution rates of all molecules changes. Since we wanted to find the regulatory degrees of freedom for the cell, the EGSs were not very useful: biological environments are noisy which means that these degrees of freedom would fluctuate constantly. That is why we defined the Elementary Growth Modes as equivalence classes of Elementary Growth States: two EGSs with the same set of expressed enzymes belong to the same EGM. Under small fluctuations in the environment, the set of EGSs changes, but the set of EGMs is constant. The set of growth-supporting EGMs *can* change due to a larger change in the environment, e.g., due to the depletion or appearance of nutrients.

The EGMs are analogous to EFMs, objects defined in models where no explicit synthesis of enzymes and the ribosome is considered. The EFMs are defined as the minimal (i.e., non-decomposable) combinations of reaction rates that support a steady-state flux. So, where the EGMs are the minimal building blocks of balanced growth, the EFMs are the building blocks of metabolism. While the EGMs are defined as equivalence classes of EGSs, the EFMs are defined up to a normalization factor, in other words, the EFMs are equivalence classes of steady-state flux vectors where vectors are identified that differ only by an overall factor. Moreover, we showed that, as long as dependent metabolites (for example due to so-called moieties) are removed, an EGM will have exactly as many active metabolic reactions as non-zero metabolites; the ribosome-catalyzed synthesis adds another reaction. EFMs, also after removing dependent metabolites, have one active reaction more than the number of used metabolites. The additional reaction in the EFM is the (virtual) biomass reaction, thus playing a similar role as the ribosome-catalyzed reaction.

### Growth rate is maximised by using a small number of active EGMs

Evolutionary theory suggests that in stable environments, those microorganisms are selected that express the phenotype that maximises the growth rate. One of our aims was therefore to describe, in any given condition, the phenotype with the maximal growth rate.

We proved in Theorem 5 that the maximally possible growth rate in any fixed environment is attained in an EGM. This makes intuitive sense: by reducing the number of active EGMs, the redundancy in the reaction network is reduced. When fewer different proteins are used, the remaining ones can be expressed to a higher level. To maximise the growth rate, the cell should thus express only the pathways that are most efficient in terms of resources. This ultimately leads to cellular states in which no redundant reactions are active, i.e., with only one active EGM, such that none of the reactions can be removed without stopping growth. Again, EGMs prove to be analogous to EFMs, since it was proven that the metabolic flux through a proteome-constrained metabolic network is maximised in an EFM (Wortel et al., 2014; Müller et al., 2014).

Because cells and their compartments have a limited size, and because each molecule takes up a certain volume, molecule concentrations in a cell are bounded. As a consequence, biochemical constraints arise for weighted sums of molecule concentrations. These give rise to constraints on the enzyme concentrations, and thus on the possible enzyme expression patterns. These constraints might change the optimal solution. For example, an EGM that is very efficient in terms of cytosolic volume might be very inefficient with respect to the limited membrane area. The first of these constraints was already incorporated when we assumed a constant osmotic pressure (Equation (6)), but additional constraints may be active and can be imposed on the model. We proved (Theorem 7) that a small number of EGMs still constitute the optimal solution when additional constraints are added. However, one EGM might be added for each constraint that is imposed. Given that the constant osmotic pressure constraint was already active, the number of active EGMs is thus bounded by the number of constraints on enzyme expression. We hereby generalized yet another existing result for EFMs to their self-fabricating counterparts (de Groot et al., 2019).

Although the above two maximiser theorems have analogs for EFMs, we needed to devise a completely different method to prove them. In EFM theory, the objective flux can be maximised by fixing this flux to a specific value after which the enzyme costs are minimized (Wortel et al., 2014; Müller et al., 2014; de Groot et al., 2019). The equivalence between the maximisation of growth rate and the minimisation of enzyme costs does not extend due to the nonlinear nature of EGMs. Indeed, in EGM theory the enzyme costs are captured by the ribosome occupation |***α***|, and if ***α*** is an EGS, this does not imply that a multiple of ***α*** is too since the balanced growth polytope changes with *μ*. This nonlinearity is also the reason that we could not compare the enzyme costs of several EGMs with the ‘cost vector formalism’ that we introduced before (de Groot et al., 2019). The absence of such a tool is certainly a disadvantage of the more general EGM-theory. Nevertheless, we have shown that an increase of the overall ribosomal capacity λ in the constraint |***α***| ≤ λ, will never lead to a decrease of the maximal growth rate. It is unclear if a more general duality exists.

The theory thus predicts that evolution of microorganisms towards higher growth rate will involve a simplification of their metabolisms towards a minimal, combined set of EGMs that gives the highest growth rate. The number of active EGMs will depend on the number of growth limiting constraints. Evolution of metabolism proceeds via mutations, either affecting the kinetics of enzymes, to increase the activity per unit enzyme, or affecting the regulation of protein expression. The first type of evolution will change the properties of the EGMs, while the second type will change the number or identity of EGMs that is expressed. The above theorems involve the second type of evolution; they predict that – no matter the environment, nor the specific properties of the growth-supporting EGMs – only a small number of EGMs should eventually be selected. How close microorganisms currently are to this optimal state is unclear, and awaiting experimental investigation, although indirect evidence is mounting that cells are very good indeed (Keren et al., 2016; de Groot et al., 2019; Planqué et al., 2018).

### EGMs are the basic building blocks of cellular growth models

In order to fully describe the phenotypes that can sustain balanced growth, we needed to incorporate 1) a direct coupling between reaction rates and growth rate, 2) explicit synthesis of enzymes and the ribosome, and 3) nonlinear and metabolite-dependent enzyme kinetics. All three aspects added nonlinearity to our theory; the first makes it impossible to prescribe the growth rate and solve for all remaining variables in steady state, the second makes the solutions depend nonlinearly on the growth rate, and the third introduces a nonlinear dependence on metabolite concentrations. In the resulting theory, all phenotypes and their corresponding growth rates can in principle be calculated, but these computations are currently not feasible for a genome-scale model. We thus developed a quantitative framework that led to general theoretical results, but no quantitative predictions can be made for a specific organism at present.

Fortunately, our theory can be simplified in various ways, resulting in computationally feasible models. Elementary Growth Modes are still the minimal building blocks, and the above-mentioned maximiser theorems still hold. A commonly used simplification is to consider a small coarsegrained whole-cell model (Scott and Hwa, 2011; Molenaar et al., 2009; Weiße et al., 2015; de Jong et al., 2017). These models become computationally feasible because of their small size, such that no further simplifications have to be made; EGM-theory is directly applicable to this type of models. Another popular modelling approach ignores the explicit synthesis of enzymes and the ribosome, replacing it by a constant biomass reaction as a sink for cell components. These approaches either consider medium-scale models with nonlinear enzyme kinetics (Wortel et al., 2018; Khodayari and Maranas, 2016), or genome-scale models without metabolite concentrations and only a maximal catalytic rate (*k*_cat_) as enzyme kinetics(Beg et al., 2007), or genome-scale models without enzyme kinetics (FBA (Orth et al., 2010)). In all these approaches, the solutions can be written as convex combinations of EFMs, which can be seen as the linear approximations of EGMs (Section 2.8). The recently developed Metabolism and Expression (ME) models, currently make the least simplifying assumptions in modelling cellular growth (Goelzer et al., 2009; O’Brien et al., 2013, 2016): enzyme and ribosome synthesis are explicitly modelled, but enzyme kinetics are replaced by a single catalytic rate. The minimal building blocks of these models are thus EGMs where the saturation function is replaced by a constant and the dilution of metabolites is ignored. Very recently, theory was also developed for kinetic models with explicit synthesis of one type of protein, but without direct coupling between the reaction rates and the growth rate (Dourado and Lercher, 2019). This simplification enabled the authors to further analyse the optimal growth state, which was indeed a simplification of an EGM.

In summary, our theory gives a very general description of the steady states of self-fabricating cells, thereby providing the theoretical background and identifying the minimal building blocks for all the commonly used simplifications of this description.

### A constant amino acid composition greatly simplifies self-fabrication

Experimental data suggests that the amino acid composition of cells is constant across different growth conditions (Figure 4). This can only be maintained when amino acids are consumed at rates with constant ratios. Under that assumption, EGMs simplify considerably, and become much more “linear” objects: the flux values of the EGM can be approximated by an EFM. Under the additional assumption that the dilution of metabolites is negligible, and within one EGM at fixed metabolite concentrations, the fractions of the ribosome allocated to producing enzymes, the *α_j_*’s, all have the same dependence on the growth rate (Theorem 10). In other words, the relative allocation of these enzymatic *α_j_*’s remains equal. The ribosomal fraction allocated to producing ribosomes, *α*_*n*+1_, does scale differently: this fraction changes linearly with the growth rate. The result of this simplification is therefore that to control enzyme/ribosome synthesis the cell should only tune two different variables: the overall fraction of the ribosome allocated to making enzymes, and the fraction of the ribosome allocated to ribosome production. The ppGpp-mechanism in *E. coli*, which acts on amino acid abundances and controls the balance between metabolism and ribosome synthesis, fits perfectly with this structure (Bosdriesz et al., 2015).

Due to this simplification, linear growth laws may still be expected, even though self-fabrication is intrinsically nonlinear. For example, since EGMs are the building blocks of the so-called Metabolism and Expression models, we expect that each EGM in the model can be well-approximated by a single EFM. Indeed, in (O’Brien et al., 2013) it is observed only in the case of sulfur- and magnesium-limited growth that “growth is sometimes non-linear as a function of uptake rate, due to changing biomass requirements”.

### Stresses and non-enzymatic reactions can be considered in terms of EGMs

Although the theory is presented for whole-cell models that are geared solely towards growth and self-fabrication, it does not exclude non-enzymatic (i.e., diffusive) reactions, stress responses, and maintenance or homeostasis activities. Processes that are not directly related to growth, such as heat shock responses or the removal of toxins, may all be incorporated. As long as the proteins involved in those processes act linearly on the reaction rates, growth rate maximisers will again be EGMs of the corresponding model. In Figure 5 we provide an example of a situation where it is optimal to invest in a stress response, in this case the removal of a toxin. The effect of the toxin was modelled by adding an inhibitory effect of the toxin on all metabolic enzymes.

**Figure 5:**
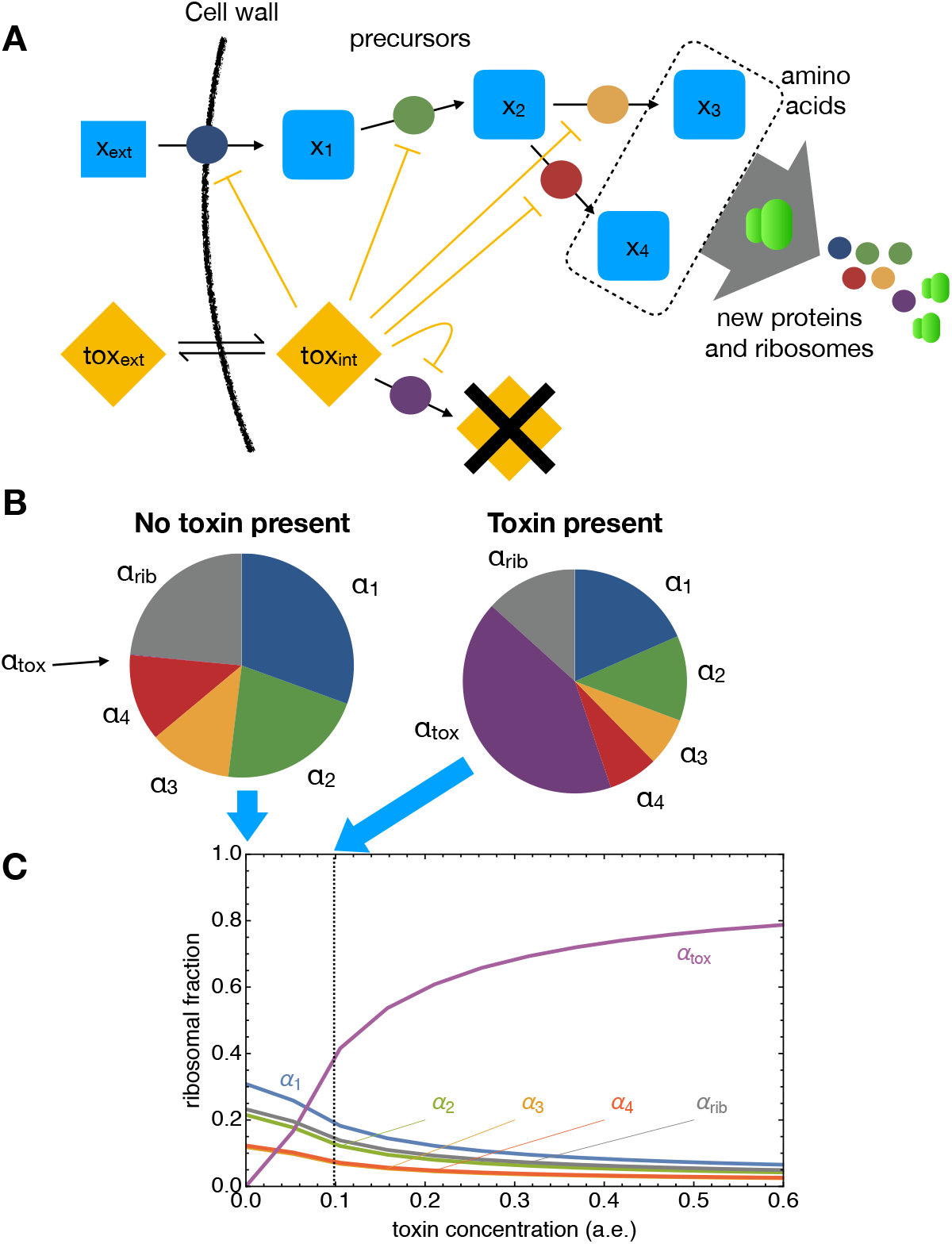
Illustration of investment in toxin removal while maximising growth rate. **(A)** The example network contains five metabolites; *x*_1_ to *x*_4_ are precursor molecules and *x*_3_ and *x*_4_ play the role of amino acids. An external toxin can diffuse over the membrane and, when inside the cell, inhibits the catalytic activity of all proteins. A fifth protein, in purple, destroys the toxin. **(B)** Without toxin present, investment is divided over the four proteins and the ribosome to maximise steady state growth rate. With toxin present, the toxin-degrading protein is also expressed, even though it does not directly contribute to growth. However, without expressing this stress protein, the maximal growth rate would have been lower. **(C)** Investment in proteins and ribosomes to attain maximal balanced growth rate as a function of toxin concentration. Note that in this example, a switch is observed, from an EGM with four metabolites to one with five. Full detail and code may be found in the SI.

### Open problems and outlook

Clearly, many open theoretical problems remain. For example, we have described the (optimal) phenotypes that lead to steady-state self-fabrication in static environments, but we do not know if the same phenotypes are evolutionary favourable in dynamic environments. Moreover, even if we know that the optimum is attained in one, or a combination of a few, EGMs, we do not know how a cell should find these optimal protein expression states with its regulatory circuitry. Further, it is unclear to us if more existing results for EFMs can be generalised to EGM theory. It is known, for example, that finding the metabolite concentrations that maximise the specific flux in an EFM is a strictly convex optimisation problem (Noor et al., 2016; Planqué et al., 2018), but we do not know if an analogous result for EGMs can be proved.

We hope that answering this type of open theoretical questions will guide us in studying microbial physiology. We believe that a sound fundamental theory could help to better understand the common denominators underlying qualitatively similar physiological behaviours of evolutionary distinct microorganisms. Systems biology is, however, remarkably short of experimentally testable theories, in contrast to other systems sciences, such as statistical physics and population genetics. This is somewhat surprising since it is apparent, even from the history of microbiology itself, that abstract theory can aid in the understanding of concrete phenomena. The understanding of single-cell physiology has profited greatly, for instance, from theories on stochastic fluctuations in molecular circuits, and the introduction of enzyme kinetics theories revolutionised enzyme biochemistry.

We hope that this paper contributes to a growing body of ‘biomathematical’ theories that eventually provide basic answers to the molecular basis of life – firstly of microorganisms. We are convinced that such a theory is within reach. The next frontier we foresee is understanding metabolic behaviour of microorganisms in terms of growth rate maximisation and constrained protein expression – a general biomathematical theory that merges enzyme biochemistry, metabolic network reconstructions and evolutionary theory.

## Supporting information

Supplementary Analyses and Matlab-Code

## Footnotes

1 The only alternative would imply that the weighted sum, 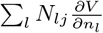, is constant in time for all chemical reactions *j* while the 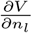 are not. This is unlikely, so we will indeed assume that 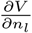 is independent of time.

2 If we assume that the osmotic pressure is kept constant in a cell, the import of a particle must lead to the import of water. The molar volume parameters, *ρ_l_*, will therefore not only comprise the volume of particle *l*, but also the volume of the water that is needed to keep the osmotic pressure constant. These parameters thus depend on the osmotic pressure.

