## Supplementary Analyses and Matlab-Code for "Elementary Growth Modes provide a molecular description of cellular self-fabrication"

April 12, 2019

### 1 Including non-enzymatic reactions

The metabolic ODEs are now augmented with non-enzymatic reactions with rate vector $\boldsymbol{u}(\boldsymbol{x})$, and stoichiometry matrix $S$, so that

$$\dot{x}_k = \sum_j S_{kj} u_j(\boldsymbol{x}) + \sum_j P_{kj} e_j f_j(\boldsymbol{x}) - \sum_j M_{kj} r \alpha_j g_j(\boldsymbol{x}) - \mu x_k.$$

Setting $c_j = \sum_k \rho_k S_{kj}$, the definition of $\mu$ may be succinctly phrased as

$$\mu = \boldsymbol{c} \cdot \boldsymbol{u} + \boldsymbol{a} \cdot \boldsymbol{v} + \boldsymbol{b} \cdot \boldsymbol{w}.$$

The steady state equations for $\boldsymbol{x}$ now read

$$S\boldsymbol{u} + P\boldsymbol{v} - M\boldsymbol{w} = \boldsymbol{x}(\boldsymbol{c} \cdot \boldsymbol{u} + \boldsymbol{a} \cdot \boldsymbol{v} + \boldsymbol{b} \cdot \boldsymbol{w}).$$

Using the same line of reasoning as before, (**??**) becomes

$$r\left( x_k \left[ \sum_{j=1}^{n} \left( \frac{a_j f_j(\boldsymbol{x})}{\mu} + b_j \right) \alpha_j(\boldsymbol{x}) + b_{n+1}\mu \right] - \sum_{j=1}^{n} \left( \frac{P_{kj} f_j(\boldsymbol{x})}{\mu} + M_{kj} \right) \alpha_j(\boldsymbol{x}) + M_{k(n+1)}\mu \right) = \sum_{j=1}^{n} (S_{kj} - x_k c_j) u_j(\boldsymbol{x}). \quad (1)$$

Note that the ribosome concentration now appears in the equations, since it occurs on the left hand side, but not on the right due to the nonenzymatic reactions.

### 2 Including a positivity constraint for the ribosome concentration

The balanced growth equations were derived by first solving the steady state ribosome and enzyme equations, and substituting the result into the steady state equations for the metabolites. Since $\dot{r} = r(\alpha_{n+1} - \mu)$, one does not get immediate information about $r$ in steady state. Since $r$ eventually drops out of the balanced growth equations, because it is a factor in all of them, one needs to derive the steady state ribosome concentration a posteriori. The expression for $r$ is given in (**??**),

$$r = \frac{\mu}{\sum_{j=1}^{n} \left( \frac{a_j f_j(\boldsymbol{x})}{\mu} + \sigma_j - b_j \right) \alpha_j + (\sigma_{n+1} - b_{n+1})\mu}.$$

Since finding any balanced growth solution generally starts with prescribing some metabolite concentration vector $\boldsymbol{x}$, it is not guaranteed that the resulting ribosome concentration is positive. If it is not, the enzyme concentrations in steady state are all negative as well, since $e_j = r\alpha_j/\mu$.

For certain choices of $\boldsymbol{x}$, and growth rate $\mu$, some vectors in the polytope $\mathcal{P}_{\boldsymbol{x},\mu}$ may correspond to negative ribosome and enzyme concentrations. These should of course be discarded. One could incorporate an additional constraint into the polytope, requiring that the denominator of (**??**) be positive,

$$\sum_{j=1}^{n} \left( \frac{a_j f_j(\boldsymbol{x})}{\mu} + \sigma_j - b_j \right) \alpha_j + (\sigma_{n+1} - b_{n+1})\mu \geq 0.$$

This constraint is again of the same nature as the ones in $B(\boldsymbol{x}, \mu) = \mu \boldsymbol{u}_{m+1}$ in the definition of $\mathcal{P}_{\boldsymbol{x},\mu}$. Choosing a metabolite vector $\boldsymbol{x}$ such that this constraint is violated for all $\boldsymbol{\alpha} \in \mathcal{P}_{\boldsymbol{x},\mu}$ implies that the polytope to which the ribosomal constraint is add would be empty.

Any biologically reasonable balanced growth solution would have a finite, strictly positive ribosome concentration. Therefore, such solutions will not hit this new constraint.

### 3 Including stress responses

Investments in stress responses are often viewed as orthogonal to investments in growth. In EGM theory, this is not necessary. For example, a heat shock response might involve increased expression of chaperone proteins that accelerate the refolding of denaturated proteins. Chaperones themselves do not contribute to the catabolic or anabolic parts of metabolism, and are therefore viewed as not contributing to growth. The chaperones, however, extend the lifetime of proteins (they lower their natural degradation rate, which we have ignored in the whole-cell model but can be put in without any modification of the main results). In this indirect way, chaperones of course increase the growth rate relative to the situation in which they are absent.

An example implementation would be as follows. First we incorporate degradation rates of enzymes,

$$\dot{e}_j = r\alpha_j - \mu e_j - d_j(T, \boldsymbol{h}) e_j$$

in which $d_j(T, \boldsymbol{h})$ are the degradation rate of enzymes, as a function of temperature (denaturation) and chaperone concentrations $\boldsymbol{h}$. Next, the chaperones need to be synthesized

$$\dot{h}_k = r\alpha_k - \mu h_k,$$

(here modelled without its own degradation rate, but one could put that in) then one would obtain a new whole cell model, with extended stoichiometry for all the enzymes and now also chaperones.

| Variable/ parameter | Description | Unit |
|---|---|---|
| $\boldsymbol{n}$ | copy number of cellular compounds | mol |
| $\boldsymbol{c}$ | concentration of cellular compounds | mol$L^{-1}$ |
| $\boldsymbol{x}$ | concentration of metabolites | mol$L^{-1}$ |
| $\boldsymbol{e}$ | concentration of enzymes | mol$L^{-1}$ |
| $r$ | concentration of ribosome | mol$L^{-1}$ |
| $\alpha_j$ | ribosomal fraction allocated to produce enzyme/ribosome | n.a. |
| $N$ | overall stoichiometric matrix | n.a. |
| $P$ | metabolic stoichiometric matrix | n.a. |
| $M$ | metabolite-to-enzyme stoichiometric matrix | n.a. |
| $I$ | identity matrix | n.a. |
| $\mu$ | growth rate | $s^{-1}$ |
| $\rho_k$ | volumetric parameter for metabolic compound $k$ | mol$^{-1}L$ |
| $\sigma_k$ | volumetric parameter for enzymes and ribosome | mol$^{-1}L$ |
| $f_j$ | enzymatic rate law | $s^{-1}$ |
| $g_j$ | enzyme synthesis rate law | $s^{-1}$ |
| $v_j$ | metabolic rate | mol$L^{-1}s^{-1}$ |
| $w_j$ | enzyme synthesis rate | mol$L^{-1}s^{-1}$ |
| $a_j$ | net contribution to growth rate by metabolic reaction $j$ | mol$^{-1}L$ |
| $b_j$ | net contribution to growth rate by enzyme/ribosome synthesis reaction $j$ | mol$^{-1}L$ |
| $\mathcal{C}_{\boldsymbol{x},\mu}$ | cone corresponding to metabolite concentration $\boldsymbol{x}$ and growth rate $\mu$ | n.a. |
| $\mathcal{P}_{\boldsymbol{x},\mu}$ | polytope corresponding to metabolite concentration $\boldsymbol{x}$ and growth rate $\mu$ | n.a. |
| $\boldsymbol{u}_j$ | $j$-th elementary unit vector | n.a. |

Table 1: Overview of all variables and parameters used in this paper.

The number of $\alpha_j$ increases: the ribosome now needs to be allocated not only over all enzymes and the ribosome, but also over the chaperones.

If the degradation depends linearly on the chaperone concentration, the resulting balanced growth equations would have the same qualitative properties as before. The polytope $\mathcal{P}_{\boldsymbol{x},\mu,T}$ of all balanced growth solutions at metabolite concentrations $\boldsymbol{x}$ with growth rate $\mu$ and temperature $T$ now has new vertices, which are again called EGMs. Depending on $T$, the maximal growth rate solution may have a positive $\alpha_k$ for the chaperones. In that case, the investment of synthesising chaperones leads to a higher growth rate than if this investment were not made.

Similar implementations may be given for other stress responses. A second example is cleaning up of toxins (see Figure **??** in the main text, and code below), either made as an inevitable byproduct of certain metabolic reactions, or passively diffuse into the cell. Such toxins could for instance lower the $k_{\text{cat}}$ of metabolic reactions, leading to a lower growth rate. As long as cleaning up such toxins using specific proteins is implemented using kinetics that are linear in those protein concentrations, the EGM theory applies. If the effect of the toxin is sufficiently detrimental to metabolic reaction rates, the growth rate maximiser will feature investment into proteins that naturalise those toxins.

### 4 Matlab code for the toxin example in Figure ??

In this code, the $\alpha_j$ are replaced by $\beta_j$, which are defined by $\beta_j = \alpha_j g_j(\boldsymbol{x})$).

```
function [toxext,xopts,alphas] = optimal_allocation_for_toxins(net);

xset=[1:6];
eset = [1:7 9];

[vars,ftns,gtns,bgc,dBGdx,algeqs,ALGEQS] = load_functions(net,0,xset,eset);

toxext = linspace(0,1,20);
f=waitbar(0);
setup=net;
for i=1:length(toxext);
  f=waitbar(i/length(toxext),f,num2str(i));
  setup.extconc(2)=toxext(i);
  net = update_network(net,setup);
  alpha0 = rand(9,1);
  alpha0 = alpha0/sum(alpha0);
  [xopt,alpha_opt] = maximise_over_alphas(net,alpha0);
  xopts(:,i) = xopt;
  alphas(:,i) = alpha_opt;
end
close(f);
opts = [xopts; alphas(1:7,:); zeros(1,20); alphas(8:9,:)];
save('optima_toxin_resistant.mat','opts');
% for plotting in Mathematica we also want toxext (external toxin concentration)
% and sorted by column, not by rows
opts = [toxext(:) opts'];
csvwrite('optima_toxin_resistant.csv',opts);

%%%%%%%%%%%%%%%%

function s = code_BG(net,xset,eset);

% function s = code_BG(net,xset,eset);
%
% this function generates the equations for balanced growth in x,
% beta (=alpha * g(x)) and mu,
% for a general reaction network
%
% fc and gc contain the code for the reaction functions, but are not
% themselves symfun(...)'s. This is because the entire BG code needs to be evaluated as
```

```
% a symfun function at the end.
%
% The BG equations are
% xk [\sum_{j=1}^n (\frac{a_j f_j(\bb x)}\mu + \sig_j - b_j) \beta_j
% + (\sig_r - b_r)\mu ] - \sum_{j=1}^n (\frac{P_{kj} f_j(\bb x)}\mu - M_{kj} )\beta_j
% + M_{kr} \mu

fc = net.fmc;
gc = net.gmc;

for i=1:length(xset)
  x{i} = strcat('x',num2str(xset(i)));
end

for i=1:length(eset)
  beta{i} = strcat('beta',num2str(eset(i)));
end
P = net.stoichmetab;
M = net.stoichenz;
sigma = net.sigma(:);
a = net.a;
b = net.b(:);
d = sigma - b;

P = P(xset,eset);
M = M(xset,[eset,length(sigma)]);
sigma = sigma([eset,length(sigma)]);
a = a(xset);
b = b([eset length(b)]);

% \sum_{j=1}^n (\frac{a_j f_j(\bb x)}\mu + \sig_j - b_j) \beta_j + (\sig_r - b_r)\mu ]

for i=1:length(xset)
  som{i} = strcat('(',num2str(a(i)),'*(',fc{eset(i)},')/mu + ...
  ',num2str(d(i)),')*', beta{i});
end

totalsom = [];
for i=1:length(som)
  totalsom = strcat(totalsom,som{i});
  if i<length(som)
    totalsom = strcat(totalsom, '+');
  end
end
totalsom=strcat(totalsom,'+',num2str(d(end)),'*mu');
```

```
for i=1:length(som)
  rhs{i} = metab_RHS(i,P,M,xset,eset,fc);
end

BGcode = '[';
for i=1:length(xset)
  BGcode = strcat(BGcode, x{i},'*(',totalsom,')',rhs{i});
  BGcode = strcat(BGcode,';');
end

% now add the last line of the BG equations, sum alpha_i = 1.
sum_alphas = [];
for i=1:length(xset),
  sum_alphas = strcat(sum_alphas,beta{i},'/(',gc{i},')+');
end
sum_alphas = strcat(sum_alphas,'mu/(',gc{end},')-1');

BGcode = strcat(BGcode,sum_alphas,']');
s = strcat('BGCODE = symfun(',BGcode,', vars)');


%%%%%%%%%%%%%

function totalcode = metab_RHS(k,P,M,xset,eset,fc);

% - \sum_{j=1}^n (\frac{P_{kj} f_j(\bb x)}\mu - M_{kj} )\beta_j  + M_{kr} \mu

N = size(P,2);

for i=1:N
  PP{i} = num2str(P(k,i));
  MM{i} = num2str(M(k,i));
  beta{i} = strcat('beta',num2str(eset(i)));
end

MM{N+1} = num2str(M(k,N+1));

for i=1:size(P,2)
  code{i} = strcat('-(',PP{i},'*(',fc{eset(i)},')/mu - ',MM{i},') * ',beta{i},'+' );
end

totalcode = [];
for i=1:size(P,2)
```

```
  totalcode = strcat(totalcode,code{i});
end
totalcode = strcat(totalcode,MM{N+1},'*mu');

%%%%%%%%%%%%%%%

function yout = expand(yin,set,n);

% function yout = expand(yin,set,n);
%
% for vector input, this function expands a vector [yin(1) ... yin(end)]
% to a vector yout of length n, such that yout(set) = yin, and zeros elsewhere.
%
% for matrix input yin, this function expands a matrix
% yout with the same row dimension but column dimension n, such that
% yout(:,set) = yin, and zeros elsewhere.

[N,M] = size(yin);
if size(yin,1) ~= length(set) && size(yin,2) ~= length(set)
disp('the row or column dimension of yin needs to be the same as the length of set')
yin
set
end
if N==1 || M==1
  yout = zeros(n,1);
  yout(set) = yin;
else
  yout = zeros(N,n);
  yout(:,set) = yin;
end

%%%%%%%%%%%%%%%%

function [t,y,mu,alpha] = fixed_alpha_system...
(net,xset,eset,Tspan,IC,alpha,newfunctions,doplot);

% function [t,y,mu] = ...
% fixed_alpha_system(net,xset,eset,Tspan,IC,alpha,newfunctions,doplot);
%
%

if newfunctions
  [vars,args,xbargs,ftns,gtns] = load_kinetics_ftns(net,xset,eset);
else
  filename = strcat(net.name,'_all.mat');
```

```
  load(filename);
end

nmetab = net.nmetab;
nenz = net.nenz;
rho = net.rho;
sigma = net.sigma;

if IC == 0
  x0= net.x;
  e0= ones(net.nreactions,1);
  r0=1;
else
  x0 = IC(1:nmetab);
  e0 = IC(nmetab+1:end-1);
  r0 = IC(end);
end

% check if the weighted concentration of all compunds equals 1. If not normalise...

sumx = rho(xset)*x0(xset)';
sumer = sigma([eset end])*[e0(eset);r0];
%rescale so that sum(net.rho.*x0(:)) + sum(net.sigma.*[e0;r0]) = 1
if sumx < 1
  fac = (1-sumx)/sumer;
  e0 = fac*e0;
  r0 = fac*r0;
else
error('metabolite concentrations exceed rho*x = 1');
end
e0(net.nonenzreactions) = 1;
IC = [x0(:);e0(:);r0];

optqssa = odeset('RelTol',1e-9,'AbsTol',1e-9);

% set alpha
if nargin < 6
  alpha = [2 3 1 4 2 3 1 3 4 1]';
else
  alpha = alpha(:);
%  alpha=ones(net.nreactions+1,1);
  alpha(net.nonenzreactions)=0;
%  alpha = alpha/sum(alpha);
  alpha(:)';
end
```

```
metsys(0,IC,alpha,ftns,gtns,net,xset,eset);

RHS = @(t,y) metsys(t,y,alpha,ftns,gtns,net,xset,eset);
[t,y] = ode15s(RHS,Tspan,IC,optqssa);

if nargin<8
  doplot=0;
end

mu = plot_fixed_alpha(t,y,net,xset,eset,alpha,newfunctions,doplot);

%%%%%%%%%%%%%%%%%%%%%%%%%

function dy = metsys(t,y,alpha,ftns,gtns,net,xset,eset);

nx = length(xset);
nr = net.nreactions;
nmetab = net.nmetab;
nenz = net.nenz;
N = net.stoichmetab;
M = net.stoichenz;
rho = net.rho;
sigma = net.sigma;
xe = net.extconc;
nonenz = net.nonenzreactions;
a = net.a;
b = net.b;

allreactions = union(eset,nonenz);

x = y(1:nx);
e = y(nx+1:nx+nr);
r = y(end);

%restrict to the EGM (or other subnet)
N = N(xset,allreactions);
M = M(xset,[allreactions end]);
rho = rho(xset);

beta_dummy = ones(size(alpha));

X = expand(x,xset,nmetab);
B = expand(beta_dummy(1:end-1),allreactions,nr+1);
```

```
f = ftns(X,B,beta_dummy(end),xe);
g = gtns(X,B,beta_dummy(end),xe);
beta = alpha(:).*g(:);

V = e(:).*f;
W = r.*beta(:);

mu =  a(:)'*V(:) + b(:)'*W(:);

dx = N*V(:) - M*W(:) - mu*x(:);
de = W(1:end-1) - mu*e(:);
dr = W(end) - mu*r;

%nonenz reactions have 'dummy enzymes' that are set to 1, and do not change value.
de(net.nonenzreactions) = 0;

dy = [dx;de;dr];


%%%%%%%%%%%%%%%%%%%%%%%
function [vars,ftns,gtns,bgc,dBGdx,algeqs,ALGEQS] = load_functions(net,loadold,xset,eset);

% create all relevant functions and algebraic equations for the adaptive control on the
% restricted network. So ftns and gtns here are only the EGM values, not all the values

filename = strcat(net.name,'_all.mat');

if loadold==0

  [vars,args,xbargs,ftns,gtns] = load_kinetics_ftns(net,xset,eset);

  eval(net.syms);
  args
  xbargs

  % the Balanced Growth equations %%%%%%
  BGcode = code_BG(net,xset,eset)
  eval(BGcode);
  BGC = matlabFunction(BGCODE);
  bgc = strcat('bgc = @(x,beta,mu,z) BGC(',args,')');
  eval(bgc);

  % now make the jacobian of BGC with respect to all x and beta (except mu) variables
  dBGdx1 = strcat('DBGDX = symfun(jacobian(BGCODE,',xbargs,'),vars)');
  eval(dBGdx1);
```

```
  dBGdx11 = matlabFunction(DBGDX);
  % the eval() is to make sure that the outcome is a double, not a rational number
  dBGdx = strcat('dBGdx = @(x,beta,mu,z) eval(DBGDX(',args,'))');
  eval(dBGdx);
  %dBGdx(net.x,net.b,1,net.xe);

  % the complete set of algebraic equations used for the adaptive control.
  % the detBG function creates the proper set of determinants of the jacobian matrix
  algeqs = @(x,beta,mu,z) [bgc(x,beta,mu,z); detBG(dBGdx,x,beta,mu,z)];
  n=length(xset);
  m=length(eset);
  ALGEQS = @(y) algeqs(expand(y(1:n),xset,net.nmetab),...
 expand(y(n+1:n+m),eset,net.nenz),...
 y(n+m+1),y(n+m+2:end));

  save(filename,'vars','ftns','gtns','bgc','dBGdx','algeqs','ALGEQS');
else
  load(filename);
end

%%%%%%%%%%%%%%

function [vars,args,xbargs,ftns,gtns] = load_kinetics_ftns(net,xset,eset);

  eval(net.syms); % declare all variables syms x1 x2 ...

  if nargin < 2
  % compute all the functions, not just some EGM subset
  % this version is used to compute all function values in maxgrowthEGM for example
  xset = [1:net.nmetab];
  eset = [1:net.nreactions];
  filename = strcat(net.name,'_all.mat');
  end

  [vars,args,xargs,xbargs] = load_vars(net,xset,eset)
  eval(vars);

  allreactions = union(eset,net.nonenzreactions);

  % FTNS %%%%%%%%%%%%%%%%%%%%%
  ftns = 'FTNS = symfun([';
  for i=1:length(allreactions)
    ftns = strcat(ftns,net.fmc{allreactions(i)});
    if i < length(allreactions)
      ftns = strcat(ftns,';');
```

```
    end
  end
  ftns = strcat(ftns,'],vars)')
  eval(ftns);
  ftns1 = matlabFunction(FTNS);
  ftns = strcat('ftns = @(x,beta,mu,z) ftns1(',args,')')
  eval(ftns);

  % GTNS %%%%%%%%%%%%%%%%%%%%
  gtns = 'GTNS = symfun([';
  for i=1:length(allreactions)
    gtns = strcat(gtns,net.gmc{allreactions(i)},';');
  end
  % don't forget to always add the ribosome synthesis rate:
  gtns = strcat(gtns,net.gmc{end});
  gtns = strcat(gtns,'],vars)');
  eval(gtns);
  gtns1 = matlabFunction(GTNS);
  gtns = strcat('gtns = @(x,beta,mu,z) gtns1(',args,')');
  eval(gtns);

  if nargin < 2
    save(filename,'vars','args','xbargs','ftns','gtns');
  end
function [Vars,args,xargs,xbargs] =  load_vars(net,xset,eset);

  eval(net.syms)

  % create vars variable from xset, eset
  Vars = 'vars =['; % this is the set of all variables on which the functions depend
  args = ''; % this is the same, but without the [ ]
  xargs = ''; betaargs = ''; % the xvars and beta vars
  for i=1:length(xset)
    Vars = strcat(Vars, ' x',num2str(xset(i)));
    args = strcat(args, 'x(',num2str(xset(i)),'),');
    xargs = strcat(xargs, ' x',num2str(xset(i)));
  end
  xargs = strcat('[',xargs);
  for i=1:length(eset)
    Vars = strcat(Vars, ' beta',num2str(eset(i)));
    args = strcat(args, 'beta(',num2str(eset(i)),')');
    betaargs = strcat(betaargs, ' beta',num2str(eset(i)));
    if i < length(eset)
      args = strcat(args,',');
    end
```

```
  end


  % collect all x and beta variables
  xbargs = strcat(xargs,betaargs,']');

  xargs = strcat(xargs,']');

  Vars = strcat(Vars, ' mu ');
  args = strcat(args,',mu');

  nxext = net.nxext;
  ve = net.vextindex;
  for i=1:nxext
      Vars = strcat(Vars, ' z',num2str(i));
      args = strcat(args,',z(',num2str(i),')');
  end

  % now add toxin to the list
  if net.toxin

 for i=1:length(net.toxin)
 if ~ismember(net.toxin(i),xset)
 Vars = [Vars,' x',num2str(net.toxin(i))];
 args = [args,',x(',num2str(net.toxin(i)),')'];
 end
 end
  end

  Vars = strcat(Vars, ']');

%%%%%%%%%%%%%%%%

function net = make_network(newfunctions,setup,x,r);

% function net = make_network(newfunctions,setup,x);
%
% make a network struct.
%
% - without additional arguments, net = make_network produces a default network
% - net = make_network(1) produces a default network with newly computed kinetics
% - net = make_network(newfunctions, setup) produces a network as defined in setup.
% setup is a struct which should contain the same field names
% (though not necessarily all of them)
%  to define a network.
```

```
% - net = make_network(newfunctions, setup,x) creates
% the same network as above, but with different
%  choice of metabolite concentrations. Setting setup=struct;
% (an empty struct) and running the above
%  command allows one to change only the network's
% metabolite concentrations. This is necessary
%  to find optimal EGMs.

if nargin == 0
newfunctions = 0;
end

net = struct;

if nargin < 2 || setup == 0
net.name = 'toxin_resistant';

% parameters for the kinetics functions

P = [2.1   2.3   2.2   1.9   1.3   2.4   2.3   1.5   100  10; % kcats
         1     1.3   1.5   1.2   0.8   1.1   0.9   0.7   1.15  4.2; % Km for x1
         1     1     1     1     1     1     2     1     1    1; % Keq
         1.2   1.3   1.1   1.4   0.9   0.8   0.7   1     0.6   5; %Km for x2
         1     1     3     5     2     2     4     5     1    2; %
         1     1     1     1     1     1     1     1     1     1; % Km for toxin
         4     1     4     4     4     5     3     3     2     1;
         1     2     4     3     4     2     3     3     3    3;
         4     1     1     2     2     0     0     2     4    2;
         3     4     4     3     3     2     1     4     3    1];

 % the enzyme synthesis params.
   Q = [1     1     1     1     1     1     1     1   2;
     1     1     1     1     1     1     1     1   1;
     1     1     1     2     1     2     2     3   3;
     3     3     3     3     2     1     4     3   4;
     1     2     5     2     2     4     5     1   5;
     1     4     1     4     3     3     3     4   6;
     4     3     4     4     5     3     3     2   2;
     1     6     3     4     2     3     3     3   3;
     4     2     2     2     2     1     2     4   4];

 Q = repmat([1;2;3;1;4;1;1],1,11);

 %toxin resistant
      N = [     1 -1  -2  0  0  0  0  0  0;
```

```
        0  2   0 -1  0 -1  0  0  0;
        0  0   1  0 -2  0 -3  0  0;
       0  0   0  3  0  0  2  0  0;
       0  0   0  0  3  2  0  0  0;
      0  0   0  0  0  0  0  1 -1];


% dim M is (nmetab, nenz + 1), because of the extra ribosome
 M = [0  0  0  0  0  0  0  0  0  0; % 10* to make enzymes more costly to make
  0  0  0  0  0  0  0  0  0  0;
  0  0  0  0  0  0  0  0  0  0;
 33 28 32 30 28 32 30  0 29 27; % v8 is not enzymatic
 29 31 31 27 29 32 30  0 28 32; % but just diffusion across membrane,
  0  0  0  0  0  0  0  0  0  0];


net.stoichmetab = N;
net.stoichenz = M;
net.metabparams = P;
net.enzparams = Q;

net.vextindex=[1 8]; % ve - which reactions are import reactions?
net.extconcindex=[1 2]; % vex - which external concentrations are
% used in these import reactions?
net.extconc = [3 0.1]; % xe - external input concentration.
% second one is toxin
net.extconcindexset = [1]; % xeset - the choice of external concentrations
%that are part of the EGM
net.nonenzreactions = [8]; %nonenzymatic reactions. in this case just the toxin import
net.aminoacids = [4,5]; % identify which metabolites play the role of 'amino acids'
  % (i.e., from which the enzymes and ribosome are made)

net.sensors = [1]; % choose which internal variables will act as sensor
net.ze0 = [3]; % external predicted value

net.toxin = [6]; % specify which substance is toxic, and causes lower kcats.

net.x = [1,.1,.1,0.1,0.1,0.1]; % give an IC for the internal metabolite concentrations
    net.r = 0.01;

net.nxext = length(net.extconc); %number of external concentrations
net.nreactions = size(N,2); % ALL reactions, enz + nonenz
net.enzreactions = setdiff([1:size(N,2)],net.nonenzreactions); %enzymatic reactions.

net.nmetab = size(N,1);
```

```
 net.nenz = length(net.enzreactions);

 net.rho = 0.01*linspace(1,1,net.nmetab);
 net.sigma = 10*[20 40 30 50 20 40 30 10 1 20 4 3 2 4 3 2 3 4];
 net.sigma = net.sigma([1:net.nreactions+1]);% also here one extra for the ribosome
 net.sigma(net.nonenzreactions) = 0;
else
  % setup the network with user-specified values
  if setup
    net = update_network(net,setup);
  end
end

% make sure that if metab concentrations were
% specifically supplied, then these should be taken
if nargin>2
net.x=x;
net.r=r;
end

%%%%%%%%

% from this point on the definitions depend on the network setup

N = net.stoichmetab;
M = net.stoichenz;
P = net.metabparams;
Q = net.enzparams;

% the a_j and b_j from EGM theory.
net.a = sum(repmat(net.rho',1,net.nreactions).*N);
net.b = net.sigma - sum(repmat(net.rho',1,net.nreactions+1).*M); % also here
%one extra for the ribosome
 % the nonenz reaction entries get b=0 automatically;

% set up a matrix specifying which metabolite acts as input or output to which reaction
net = set_reactions_metab_dependence(net);

% generate kinetics
[net.fm,net.fmc,net.gm,net.gmc] = kinetics(net);

net = make_symbolic_variables(net);

if newfunctions
   [net.vars,net.args,net.xbargs,net.allftns,net.allgtns] = load_kinetics_ftns(net);
```

```
else
filename = strcat(net.name,'_all.mat');
load(filename);
net.allftns = ftns;
net.allgtns = gtns;
end

net.f = net.allftns(net.x,net.b,1,net.extconc);
net.g = net.allgtns(net.x,net.b,1,net.extconc);
function [xopt,alpha_opt] = maximise_over_alphas(net,alpha0);

xset=[1:6];
eset = [1:7 9];

F = @(alpha) -calculate_growth_rate(net,xset,eset,alpha);
A = -eye(length(alpha0)+1,length(alpha0));
A(end,:) = 1;
B= zeros(length(alpha0)+1,1);
B(end)=1;
alpha_opt = fmincon(F,alpha0,A,B);
[t,y,mu] = fixed_alpha_system(net,xset,eset,...
[0,1e6],0,[alpha_opt(1:7); 0; alpha_opt(8:9)],0,0);
xopt = y(end,xset);

%%%%%%%%%%%%

function muend = calculate_growth_rate(net,xset,eset,alpha);

[t,y,mu] = fixed_alpha_system(net,xset,eset,[0,1e6],0,[alpha(1:7); 0; alpha(8:9)],0,0);
muend = mu(end);
[alpha' muend];
sum(alpha);


%%%%%%%%%%%%%%%%

function [mu] = plot_fixed_alpha(t,y,net,xset,eset,alpha,newfunctions,doplot);

nx = length(xset);
nmetab = net.nmetab;
nenz = net.nenz;
N = net.stoichmetab;
M = net.stoichenz;
rho = net.rho;
xe = net.extconc;
```

```
nonenz = net.nonenzreactions;
allreactions = union(eset,nonenz);
ne = length(allreactions);
a = net.a;
b = net.b;

%restrict to the EGM (or other subnet)
N = N(xset,allreactions);
M = M(xset,[allreactions end]);
rho = rho(xset);

if newfunctions
  [vars,args,xbargs,ftns,gtns] = load_kinetics_ftns(net,xset,eset);
else
  filename = strcat(net.name,'_all.mat');
  load(filename);
end
x = y(:,1:nx);
e = y(:,nx+1:nx+ne);
r = y(:,end);

alpha_dummy = ones(net.nreactions+1,1);

%compute f(x) and g(x) values
beta_dummy = ones(size(alpha));

for i=1:length(t)
  X = expand(x(i,:),xset,nmetab);
  f(i,:) = ftns(X,alpha_dummy(1:end-1),beta_dummy(end),xe);
  g(i,:) = gtns(X,alpha_dummy(1:end-1),beta_dummy(end),xe);

  beta(i,:) = alpha'.*g(i,:);

  V = e(i,:).*f(i,:);
  W = r(i).*beta(i,:);
  %compute growth rates
  mu(i) = a(:)'*V(:) + b(:)'*W(:);
end

if doplot==0
  return
end

figure(1)
clf
```

```
%%%% metabolite levels

subplot(2,3,1)
hold on
Lg = strcat('legend([px],');
q = char(39); % the "'" quote
for i=1:nx
  px(i) = plot(t,x(:,i),'LineWidth',2);
  Lg = strcat(Lg,q,'x',num2str(xset(i)),q);
  if i < nx
    Lg = strcat(Lg,',');
  end
end
  Lg = strcat(Lg,');');
eval(Lg);
hold off
title('metabolite concentrations')

%%% enzyme levels

subplot(2,3,2)
hold on
Lg = strcat('legend([pe],');
q = char(39); % the "'" quote
for i=1:length(eset)
  pe(i) = plot(t,e(:,eset(i)),'LineWidth',2);
  Lg = strcat(Lg,q,'e',num2str(eset(i)),q,',');
end
pe(end+1) = plot(t,r,'LineWidth',2);
Lg = strcat(Lg,q,'Rib',q,');');
eval(Lg);
hold off
title('enzyme/ribo concentrations')

%%% enz saturation levels (incl kcats)

subplot(2,3,4)
hold on
Lg = strcat('legend([pf],');
q = char(39); % the "'" quote
for i=1:length(eset)
  pf(i) = plot(t,f(:,eset(i)),'LineWidth',2);
  Lg = strcat(Lg,q,'f',num2str(eset(i)),q);
  if i < length(eset)
```

```
    Lg = strcat(Lg,',');
  end
end
Lg = strcat(Lg,');');
eval(Lg);
hold off
title('enzyme saturations')

%%% enzyme synth rates

subplot(2,3,5)
hold on
Lg = strcat('legend([pg],');
q = char(39); % the "'" quote
for i=1:length(eset)
  pg(i) = plot(t,g(:,eset(i)),'LineWidth',2);
  Lg = strcat(Lg,q,'g',num2str(eset(i)),q,',');
end
pg(end+1) = plot(t,g(:,end),'LineWidth',2);
Lg = strcat(Lg,q,'gr',q,');');
eval(Lg);
hold off
title('enzyme/ribo synthesis')

%%% growth rate %%%%%%%

subplot(2,3,3)
hold on
plot(t,mu,'LineWidth',2)
hold off
title('growth rate')

%%% metabolic fluxes %%%%%%%

subplot(2,3,6)
hold on
Lg = strcat('legend([pf],');
q = char(39); % the "'" quote
for i=1:length(eset)
  pf(i) = plot(t,e(:,i).*f(:,i),'LineWidth',2);
  Lg = strcat(Lg,q,'v',num2str(eset(i)),q);
  if i < length(eset)
    Lg = strcat(Lg,',');
  end
end
```

```
Lg = strcat(Lg,');');
eval(Lg);
hold off
title('metabolic fluxes')
function net = update_network(net,setup);

names = fieldnames(setup);
for i=1:length(names)
value = getfield(setup,names{i});
net = setfield(net,names{i},value);
end
```
